# Molecular bases for strong phenotypic effects of single-site synonymous substitutions in the *E.coli ccdB* toxin gene

**DOI:** 10.1101/2023.03.13.532503

**Authors:** Priyanka Bajaj, Munmun Bhasin, Raghavan Varadarajan

## Abstract

Single-site synonymous mutations typically have only minor or no effects on gene function. Here, we estimate the effects on cell growth of ~200 single-site synonymous mutations in an operonic context by mutating each position of *ccdB*, the 101-residue long cytotoxin of the *ccdAB* Toxin-Antitoxin (TA) operon to all possible degenerate codons. Phenotypes were assayed by transforming the mutant library into CcdB sensitive and resistant *E. coli* strains, isolating plasmid pools, and subjecting them to deep sequencing. Since autoregulation is a hallmark of TA operons, phenotypes obtained for *ccdB* synonymous mutants after transformation in a RelE toxin reporter strain followed by deep sequencing provided information on the amount of CcdAB complex formed. Synonymous mutations in the N-terminal region involved in translation initiation showed the strongest non-neutral phenotypic effects. We observe an interplay of numerous factors, namely, location of the codon, codon usage, t-RNA abundance, formation of anti-Shine Dalgarno sequences, transcript secondary structure, and evolutionary conservation in determining phenotypic effects of *ccdB* synonymous mutations. Incorporation of an N-terminal, hyperactive synonymous mutation, in the background of the single-site synonymous mutant library sufficiently increased translation initiation, such that mutational effects on either folding or termination of translation became more apparent. Introduction of putative pause sites not only affects the translational rate but might also alter the folding kinetics of the protein *in vivo*. This information is useful in optimizing heterologous gene expression in *E. coli* and understanding the molecular bases for alteration in gene expression that arise due to synonymous mutations.

**SIGNIFICANCE STATEMENT:** Synonymous mutations which do not change amino-acid identity, typically have only minor or no effects on gene function. Using sensitive genetic screens in the context of the *ccdAB* bacterial toxin-antitoxin operon, we demonstrate that many single-site synonymous mutations of the *ccdB* toxin gene display significant phenotypic effects in an operonic context. The largest effects were seen for synonymous mutations in the N-terminal region involved in translation initiation. Synonymous mutations that affected either folding or translation termination were also identified. Lack of translational pausing due to synonymous mutations in hydrophobic residue stretches, was found to decrease the amount of properly folded CcdB protein. In summary the study provides novel insights into diverse mechanisms by which synonymous mutations modulate gene function.

## INTRODUCTION

Synonymous codon substitutions once thought to be genomic background noise, have now been widely acknowledged to have the capacity to alter protein expression, conformation and function (1). However, it is still generally thought that single synonymous mutations that preserve the identity of the amino acid and do not alter the resulting protein sequence should have minimal or no effects on cellular function or organismal fitness. Nevertheless, in most sequenced genomes, synonymous codons are used with different frequencies (2). Codon usage bias is different for different organisms, varies across different genes and also specific loci between genes (3, 4). Rare codons are often found at the N-terminal regions of ORFs in prokaryotes and eukaryotes (5). One hypothesis that is consistent with this observation is that rare codons act as a ‘ramp’ to reduce translational velocity at the beginning of the gene, also known as the ‘ramp hypothesis’ (6). Several studies suggest that poorly adapted codons at the N-terminus slow ribosome progression to increase the translational efficiency of the gene (6, 7). Many studies also report that decreased mRNA structure at the N-terminus increases gene expression (8), however there have been conflicting studies as well (9).

Protein synthesis by ribosomes takes place at non-uniform rates on mRNA (10). Synonymous mutations are known to alter gene expression by changing the translation rate via varied mechanisms such as codon usage bias (3), t-RNA abundance (11), and generation of an anti-Shine Dalgarno (aSD) sequence within the gene leading to ribosomal stalling (10). Other ways include change in the mRNA structure due to alteration in base pairing (12), or altering the mRNA steady-state levels due to either change in mRNA synthesis or degradation levels (13). However, typically multiple synonymous mutations are required for observable phenotypic effects.

In principle, translation is regulated at three different stages, initiation, elongation and termination. Translation initiation is a crucial step in protein biogenesis (14). Finding the open reading frame (ORF) and ribosome loading on the mRNA takes place at the initiation step which is the rate-limiting step and largely controls the frequency of translation of a certain mRNA (15, 16). The translation efficiency of an mRNA, i.e., the amount of protein produced per unit mRNA is primarily determined by the accessibility of the ribosome binding site, nature of the start codon, occurrence of A/U rich codons disfavouring mRNA secondary structure in the beginning of the coding gene, position of the Shine-Dalgarno (SD) sequence relative to the start codon and its complementarity to 16S rRNA (15, 17, 18). A growing body of evidence suggests that not only initiation, but also elongation plays a predominant role in controlling translation efficiency of the corresponding protein. During elongation, amino acids are added to the nascent chain one at a time. The elongation rate is non-uniform with periods of rapid movement separated by pauses (16, 19–21). Translation may also be associated with a step-by-step folding process in which partial domain folding events may be required to ensure correct folding of the entire protein (22). It is thought that some synonymous mutations can affect protein folding by affecting or targeting cotranslational folding processes (23, 24) that are altered by transient ribosome pausing (25). Alterations in translation termination may be affected by the RNA structure at the end of the coding region. Translation termination can contribute to the production of altered protein isoforms by extending the C-terminal end due to translational read-through of a stop codon (26). It is thus evident that translation efficiency of a gene is governed by the initiation, elongation and termination phases (27), but determining the relative contribution of each phase to protein abundance continues to be a challenging task.

In the context of operons, since there are two or more genes that are being translated, this adds an additional layer of complexity, as mutations can selectively affect translation efficiency of some genes of the operon relative to others (28, 29). We used the *ccdAB* toxin-antitoxin (TA) operon as a model system to infer the effects of synonymous mutations on the expression and associated phenotypes of a toxin gene, that lead to altered fitness of the organism (30, 31). The *ccd* operon from F-plasmid contains two genes, *ccdA* and *ccdB*, which encode for the homodimeric, labile antitoxin CcdA and the homodimeric stable toxin CcdB, respectively. The two proteins form a stable complex which in turn binds to the cognate *ccd* promoter and represses transcription. However, under conditions of cellular stress or plasmid loss, the labile CcdA antitoxin is degraded and CcdB binds to its cellular target, DNA Gyrase, compromising DNA replication and ultimately leading to cell death (Figure S1). Since, autoregulation is a hallmark of many TA operons, the efficiency of complex formation determines whether the operon is being repressed or derepressed, which in turn dictates its *in vivo* transcriptional levels (30). CcdB mutants can affect binding to CcdA, thereby altering CcdA-CcdB operonic regulation (32). Either altered toxin:antitoxin ratio or structural changes exhibited by CcdB can modulate CcdB expression in cells (31, 33). In the present work, we measure the fitness effects of single-site synonymous mutations spread throughout the entire *ccdB* gene. We attempt to address important issues such as 1) the relative codon-specific contribution to protein abundance for the initiation, elongation, and termination phases, 2) identification of the location of synonymous mutations that exhibit the largest phenotypic and codon-specific effects on protein synthesis, 3) understanding the molecular bases behind the observed phenotypic effects, 4) phenotypic effects of increasing translation initiation through addition of an N-terminal synonymous mutation to the existing single-site synonymous mutant library.

## RESULTS

### Phenotypes of single-site synonymous mutant library members of *ccdB* in its operonic context

A single-site synonymous mutant library of the globular cytotoxin gene, *ccdB* was made in the *ccd* operon. The *ccd* operon was cloned in pUC57 vector, a high copy number vector in order to get an amplified phenotypic response to distinguish mutant phenotype from WT (32). Each position of *ccdB* was mutated to all possible degenerate codons via inverse PCR methodology (34). All mutants were placed in identical regulatory contexts. The pooled synonymous mutant library was transformed in the CcdB resistant strain, Top10Gyr. The DNA recovered from this library was further transformed in the CcdB sensitive strain and RelE reporter strain, the latter strain is resistant to the action of CcdB and harbours a RelE reporter gene downstream of the *ccd* promoter containing the consensus Shine Dalgarno (SD) sequence (32). Following transformation and plating, DNA was recovered from pooled transformants and subsequently deep sequenced. The fractional representation of each mutant in each condition was estimated and a good correlation between the two biological replicates of the resistant strain (r = 0.99), sensitive strain (r = 0.97), and RelE reporter stain (r = 0.99) was observed when a threshold of a minimum of a 20 reads was taken in both replicates of the resistant strain as described previously (35) (Figure 1 A). Of 225 possible synonymous mutants, information for ~200 CcdB mutants were available in the resistant strain. Each synonymous mutant was assigned two variant scores, namely, Enrichment Score^CcdB^ (ES^CcdB^) and Enrichment Score^RelE^ (ES^RelE^), based on their phenotypic activity, i.e., cell growth versus cell death, which in turn is based on CcdB toxicity in the sensitive strain and RelE toxicity in the RelE reporter strain, respectively (Table S1) (see methods) (32). ES^CcdB^ scores reflect free toxin protein levels in the cell. Higher levels of free toxin result in decreased cell growth. Based on a k-means clustering algorithm, we classified synonymous mutations as hyperactive if ES^CcdB^<0.7, i.e., with a killing efficiency significantly higher than the WT and inactive if ES^CcdB^>1.8, i.e., with a killing efficiency significantly lower than the WT (Figure S2). Throughout the manuscript we assume that *ccdA* translational efficiency is unaffected by synonymous mutations in *ccdB* and that [CcdA]_TOT_ is proportional to the amount of *ccdAB* mRNA. This assumption can be justified from other studies which showed that synonymous mutations in a gene do not affect the expression levels of the upstream reporter gene in an operon (36) and that the secondary structures of mRNA in adjacent ORFs are independent of each other (28). Therefore, any change in ES^CcdB^ directly reflects a change in translational efficiency of the *ccdB* gene as we define the translational efficiency as the amount of functional CcdB produced per mRNA per unit time. Phenotypes for 15 synonymous *ccdB* mutants inferred from deep sequencing were validated in a low-throughput manner by spotting culture dilutions of each mutant in both the resistant and the sensitive strains. A good correlation of r = 0.96 was observed between the ES^CcdB^ scores and normalised CFU of mutants relative to WT, thereby validating the deep sequencing results (Figure 1 B).

**Figure 1:**
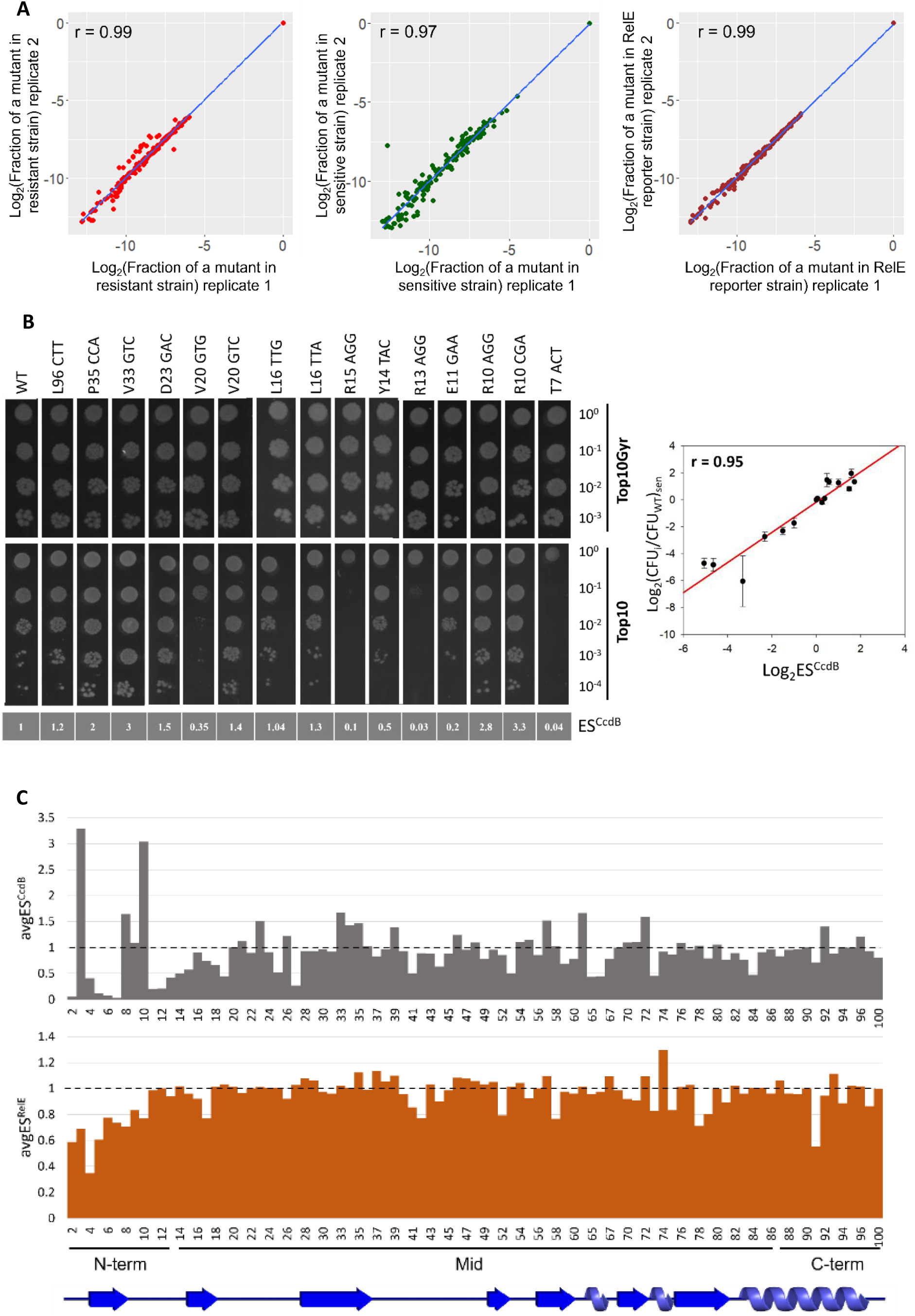
Phenotypic effects of single site-synonymous mutants of CcdB in an operonic context. (A) Correlation between biological replicates. Following deep sequencing, the fraction of each mutant normalised to the WT value is estimated for all mutants in resistant (extreme left panel), sensitive (middle panel) and RelE reporter strains (extreme right panel). (B) *In vivo* activity of CcdB mutants in both resistant (Top10Gyr) and sensitive (Top10) strains validate deep sequencing results in a low-throughput manner (left panel). Corresponding ES^CcdB^ scores of individual mutants obtained from deep sequencing are mentioned in grey in the bottom panel. The mutants are ordered according to the residue number from right to left. A correlation of r = 0.96 (*P*<.001) is observed between the ES^CcdB^ scores and normalised CFU count of mutants relative to WT in the sensitive (Top10) strain obtained in the spotting assay. Error bars for duplicate experiments are also shown (right panel). (C) Effects of synonymous mutations on ES^CcdB^ and ES^RelE^ scores as a function of location within the gene. Top panel shows position averaged ES^CcdB^ scores for synonymous mutants as a function of residue number. The bottom panel shows position averaged ES^RelE^ scores for synonymous mutants as a function of residue number. The N-terminal region shows the largest variation. Black dashed line represents the WT score. The entire data is divided into N-terminal (1-13), middle (14-86) and C-terminal (87-101) region. The secondary structure of CcdB (from PDB id 3VUB) is shown at the bottom.

To measure how efficiently the CcdAB complex is formed, the single-site synonymous mutant library was also transformed in the RelE reporter strain from which the mutants were screened based on RelE toxicity, indicated by their ES^RelE^ scores (32) (see Methods). The ES^RelE^ score for WT is 1 and due to the low dynamic range of ES^RelE^ scores, we classify mutants with ES^RelE^>1 as having a repressing phenotype and ES^RelE^<1 as having a derepressing phenotype (32). The ES^RelE^ scores combined with the ES^CcdB^ scores provide insights into the molecular mechanisms responsible for the observed phenotypes for the synonymous mutations in the sensitive strain (Table 1). Based on these two screens, all synonymous mutations are classified into four mutational categories, namely, 1) Hyperactive and Derepressing denoted as ‘H+D’; 2) Hyperactive and Repressing denoted as ‘H+R’; 3) Inactive and Derepressing denoted as ‘I+D’; and 4) Inactive and Repressing denoted as ‘I+R’, throughout the text.

**Table 1:** Possible molecular mechanisms for the observed phenotypes.

| Phenotypic category | Notation | Parameter values | Inference |
| --- | --- | --- | --- |
| Inactive and Derepressing | I+D | $ES^{CcdB} > 1.8$ and $ES^{RelE} < 1$ | Misfolded protein |
| Inactive and Repressing | I+R | $ES^{CcdB} > 1.8$ and $ES^{RelE} > 1$ | Reduced translational efficiency |
| Hyperactive and Derepressing | H+D | $ES^{CcdB} < 0.7$ and $ES^{RelE} < 1$ | Well folded, increased translational efficiency |
| Hyperactive and Repressing | H+R | $ES^{CcdB} < 0.7$ and $ES^{RelE} > 1$ | Faster folding kinetics *in vivo* or slightly increased translational efficiency |

### Synonymous mutations in the N-terminal region of CcdB display large and diverse phenotypic effects

While most synonymous mutants showed a near neutral phenotype, i.e., similar to the WT, with ES^CcdB^ and ES^RelE^ scores close to 1, a significant number show altered phenotypes (Figure 1 C). Ribosome profiling studies show that ribosomes cover 20-30 bases at a stretch (37). We examined codon specific contributions to protein abundance from three different parts of the gene, i.e., the N-terminal (residues 1-13), middle (residues 14-86) and the C-terminal (residues 87-100) regions of the *ccdB* gene. Values of ES^CcdB^ and ES^RelE^ averaged over synonymous mutations for each position were plotted to understand the overall trend. The most diverse ES^CcdB^ phenotypes relative to WT are displayed by the mutants of N-terminal region residues. The middle region shows the second highest variation in phenotypic effects. C-terminal amino acids show the least diverse phenotypic effects both in the context of CcdB toxicity as well as RelE toxicity (Figure 1 C).

### Estimating the importance of codon usage for N-terminal, middle and C-terminal region

We next examined correlations of various codon usage parameters, such as ΔGC content, RCU, CAI, RtrnaA, tAI and RCU_dV_ with ES^CcdB^, ES^RelE^, ES^CcdB^ and ES^RelE^_dV_ (see Methods section for parameter descriptions) for the entire dataset (Figure S3 A). As expected, different codon usage parameters (RCU, CAI, RtrnaA and tAI) are well correlated with each other. A positive correlation between ES^CcdB^ and codon usage parameters show that in general, enhancement in translational efficiency of CcdB is associated with decreased codon optimality (Figure S3 A). To further understand how codon bias (RCU) impacts protein abundance (ES^CcdB^) at different locations in the gene, we plotted a moving average of the two parameters over a sliding window of 5 mutants (Figure 2 A). The data showed a clear trend, i.e., ES^CcdB^ increased with increased RCU for N-terminal residues, indicating mutations to more frequently used codons decreased CcdB levels in the cell. However, for middle and C-terminal residues, this was not the case. Effects of codon usage are largest at the N-terminus, compared to the rest of the gene (Figure 2 A).

**Figure 2:**
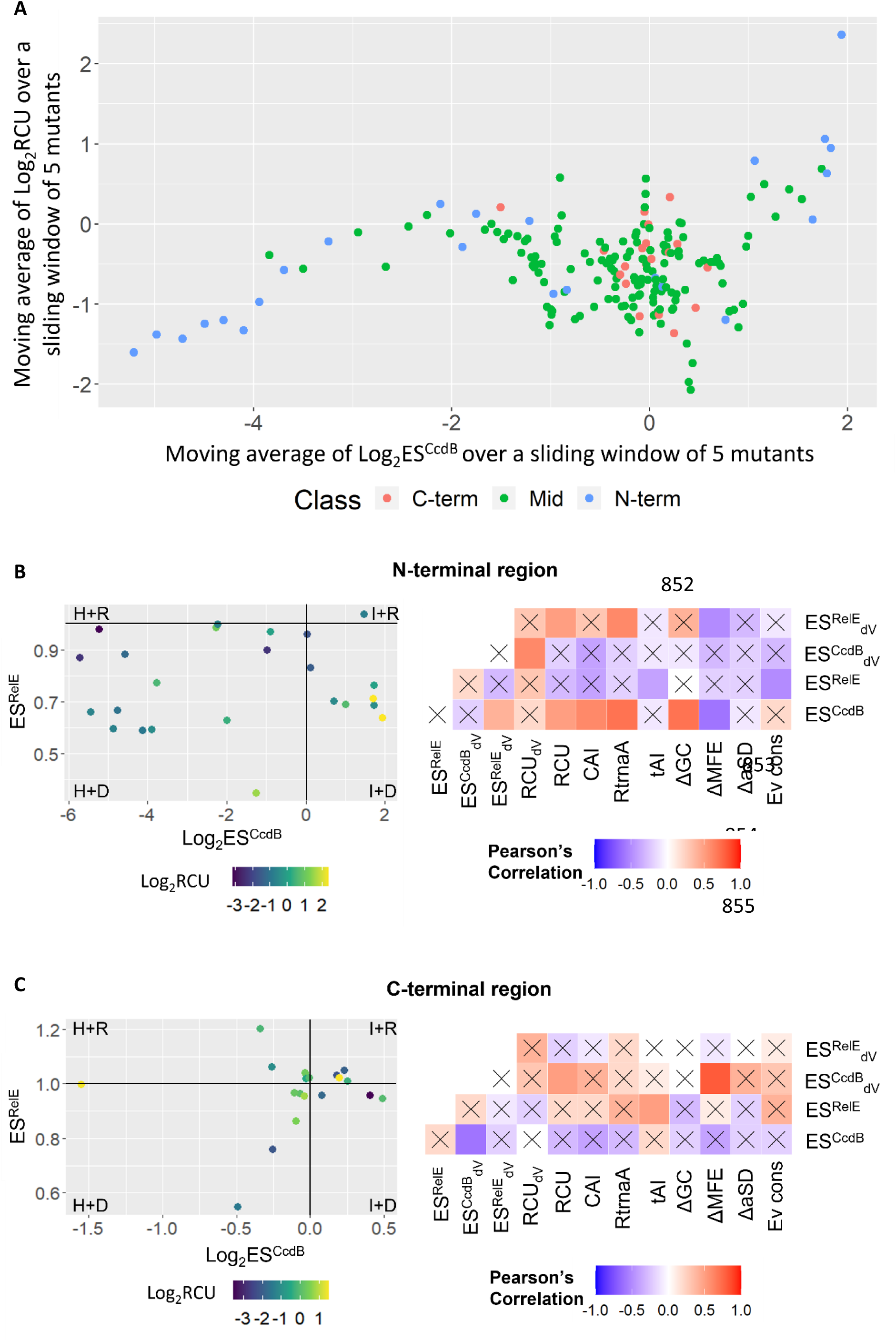
Synonymous codon usage plays an important role in regulating gene expression. (A) Altered CcdB levels (ES^CcdB^) as a function of change in codon usage (RCU) relative to WT upon synonymous substitution. Mutants in N-terminal, middle and C-terminal regions are shown in blue, green and pink, respectively. (B-C) Left panel shows the corresponding values of ES^CcdB^ (x-axis), ES^RelE^ (y-axis) and RCU (color bar) for each mutant of the (B) N-terminal region and (C) C-terminal region. WT score is represented by black lines. Right panel shows the correlation between experimentally determined relative fitness values with various sequence-based parameters for the (B) N-terminal region and (C) C-terminal region. (B-C) The right panel shows the Pearson’s correlation coefficient values and their corresponding *P* values. Gradation of white to red colour reflects positive correlation whereas the gradation of white to blue color reflects negative correlation. Non-significant correlations, i.e., correlations with *P*>0.05 are shown with a cross ‘X’ mark. Parameter definitions are in the Materials and Methods section.

63% (15/24) of the N-terminal region mutants belong to the ‘H+D’ class. Amongst these, synonymous mutations to rarer codons typically display extremely low ES^CcdB^ and ES^RelE^ scores. 21% (5/24) of the N-terminal region mutants belong to the ‘I+D’ class. Several of these synonymous mutations are at the R10 residue. Thus in this case, it appears that a synonymous mutation from a rarer to a more frequent codon might result in misfolding of the protein *in vivo* (Figure 2 B). In the N-terminal region, a positive correlation of ES^CcdB^ with RCU (r = 0.54, *P* < .01), CAI (r = 0.64, *P* < .001) and RtrnaA (r = 0.71, *P* < .001), as well as ES^CcdB^ with RCU_dV_ (r = 0.61, *P* < .001), implies that introduction of rarer codon mutants at the N-terminus increases *in vivo* translational efficiency of the protein (Figure 2 B). In the present case, a positive correlation of ΔaSD with tAI (r = 0.46, *P* < .05) and a negative correlation with RCU_dV_ (r = −0.5, *P* < .01) (Figure S3 B), indicate that introduction of rarer codons enhances the likelihood of formation of an aSD sequence, that in turn leads to stalling of the ribosome at the N-terminus. The data also might indicate that translational efficiency increases when the ribosome encounters a codon which has a significantly different codon usage than the WT. It will be interesting to see if this is a general feature across all genes.

In the middle region, ES^CcdB^ and ES^RelE^ show no correlation with codon usage parameters suggesting that alterations in synonymous codon bias in the middle region does not contribute to altering the translation efficiency of CcdB (Figure S3 C).

In the C-terminal region, most mutants lie either in the ‘H+D’ or ‘I+R’ categories, implying that the phenotypes are primarily determined by translational efficiency (Figure 2 C). ES^CcdB^ is negatively correlated to codon usage (RCU, r = −0.34, *P* = .16; CAI, r = −0.33, *P* = .06), indicating that in contrast to the N-terminus, optimal codons at the C-terminus enhance translation efficiency of the gene *in vivo*, thereby increasing its protein abundance (Figure 2 C, Figure S3 D).

### mRNA secondary structure dictates translation initiation

We next explored how predicted mRNA stability is correlated with observed phenotypes. In the N-terminal region, a negative correlation of ΔGC with ΔMFE (r = −0.61, *P* < .001) indicates that as expected, higher GC content is associated with increased mRNA stability (Figure S3 B). ES^CcdB^ is positively correlated with ΔGC (r = 0.7, *P* < .001) and negatively correlated with ΔMFE (r = −0.59, *P* < .001) (Figure 2 B), suggesting that stable mRNA might reduce translation initiation, for example by occluding the RBS, whereas absence of structure might enhance RBS binding to the anti-SD sequence of the ribosome (3, 38) or alternatively, the start codon may be more efficiently recognised by the initiator t-RNA (8).

### Evolutionary pressure drives codon selection

We also estimated evolutionary conservation for the WT *ccdB* gene both at the nucleotide and residue level. We found 61% (62/101) of the WT *ccdB* gene codons were the most conserved codons, 10% (10/101) were the least conserved codons, while the remaining 29% (29/101) have an intermediate conservation level (Figure 3 A). We further evaluated the evolutionary conservation levels of the synonymous mutant codons and compared it with their fitness effects. Largely, synonymous mutations to the most evolutionarily conserved synonymous codon display a hyperactive phenotype (Figure 3 B), indicating that synonymous codons are under selection pressure and that these mutant codons were not selected during the course of evolution for in the context of the *E.*coli CcdB gene because they will cause an increase in toxin levels leading to cell death.

**Figure 3:**
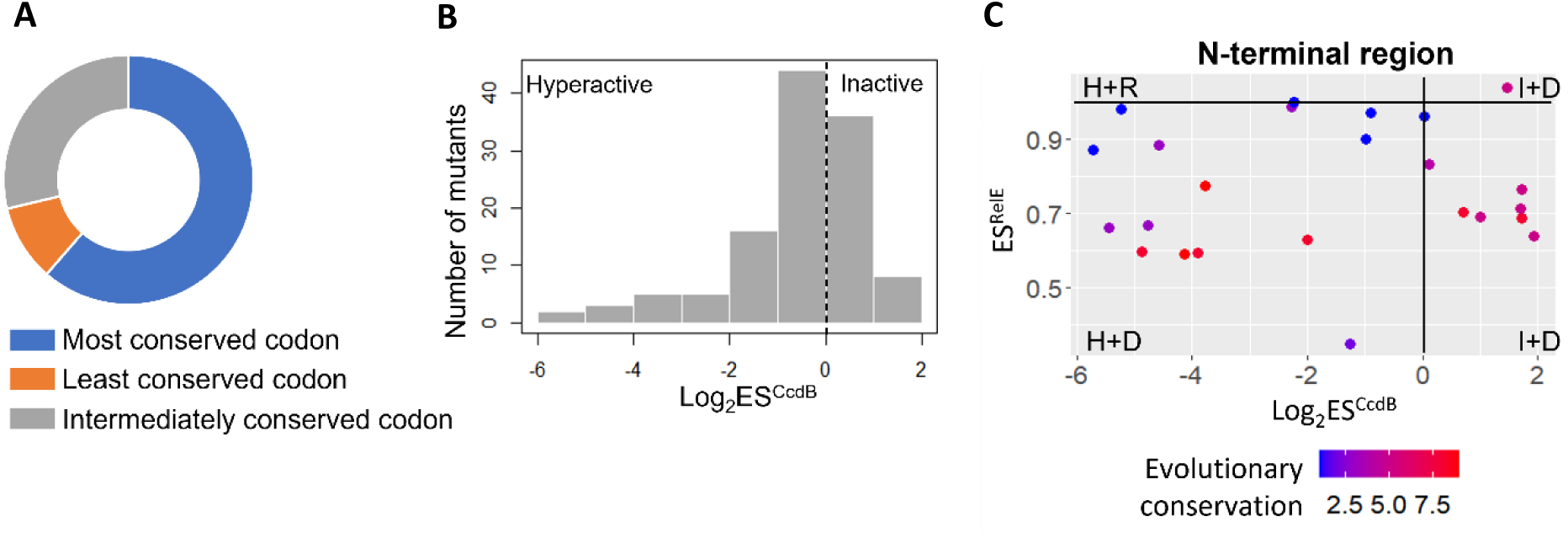
Codon selection is under evolutionary pressure. (A) Percent distribution of WT codons in context of evolutionary conservation at the nucleotide level. (B) Some CcdB positions do not have the WT codon as the most conserved codon. Instead one of the synonymous counterparts is the most conserved codon. Histogram represents the frequency distribution of ES^CcdB^ scores for synonmous mutants with the most conserved codon for the corresponding WT amino acid. (C) Corresponding values of ES^CcdB^ (x-axis), ES^RelE^ (y-axis) and evolutionary conservation at the residue level (color bar) for each mutant of the N-terminal region is plotted. Evolutionary conservation score is proportional to the conservation level of the residue.

We also probed the role of evolutionary conservation at the residue level (each residue is assigned a conservation score). Interestingly, synonymous mutations at conserved residues in the N-terminal region are enriched in either the ‘H+D’ or ‘I+D’ categories, implying that as mentioned above, these mutations either increase *ccdB* translational efficiency and CcdB protein levels, or result in a folding defect in the protein, respectively (Figure 3 C).

### Characteristics of synonymous mutations that enhance translational efficiency

The ‘H+D’ class (ES^CcdB^<0.7, ES^RelE^<1) of mutations in this study are associated with an increase in the [CcdB]_TOT_/[CcdA]_TOT_ ratio. These synonymous mutants likely result in enhanced translational efficiency of *ccdB*. Synonymous mutations in the N-terminal region, especially the 5-13 residue stretch showed the lowest ES^CcdB^ scores, implying these synonymous mutations result in the maximum fold increase in translational efficiency. Altering the synonymous codon at a single-site in the N-terminal region (R13_AGA) resulted in an ~100-fold decrease in ES^CcdB^ (Figure 4 A).

**Figure 4:**
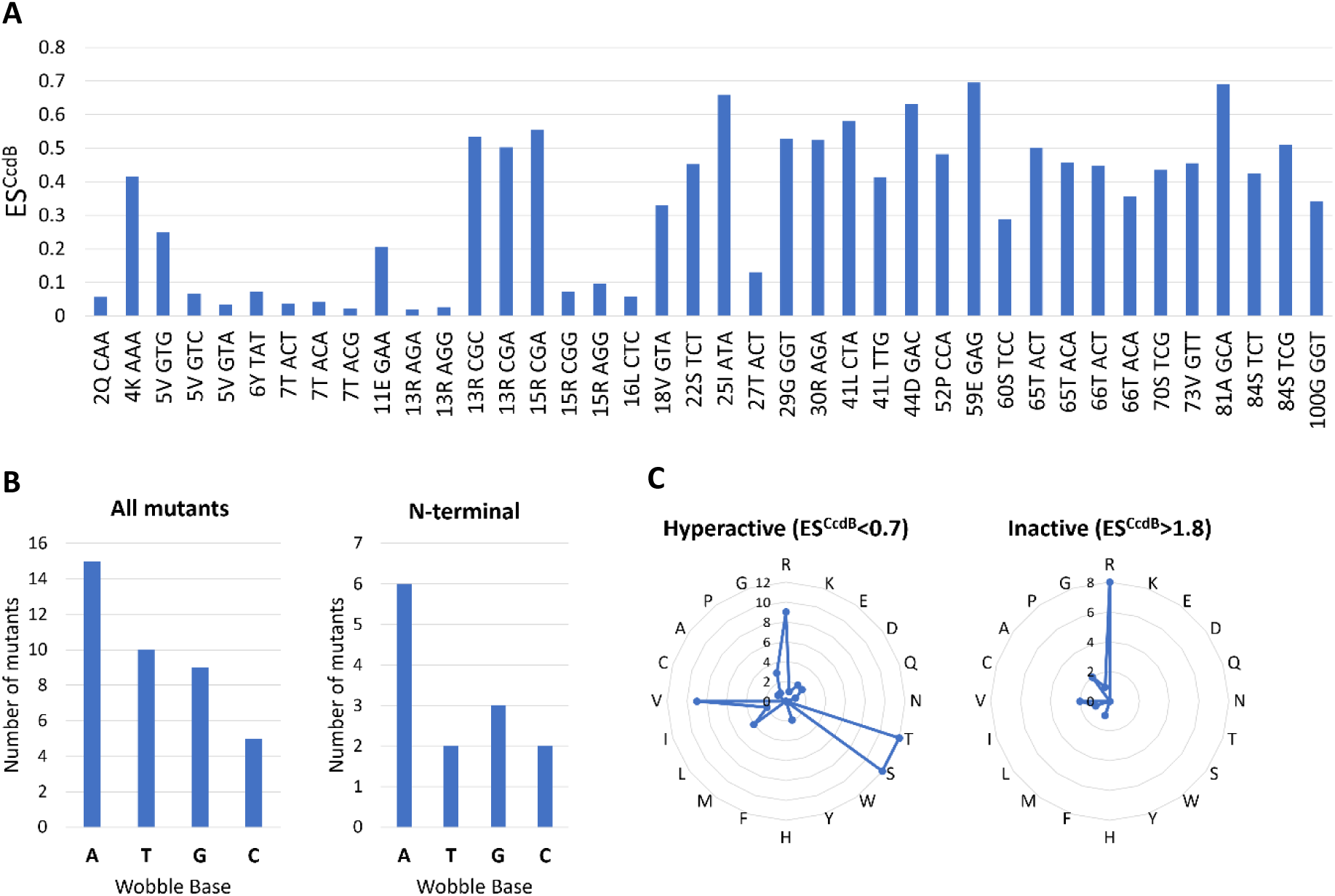
Synonymous mutations enhancing translational efficiency of the *ccdB* gene. (A) ES^CcdB^ values of synonymous mutants belonging to ‘H+D’ class (ES^CcdB^<0.7 and ES^RelE^<1). (B) Enrichment of A/T at the wobble base for all synonymous mutants belonging to the ‘H+D’ class (left panel) and for the subset lying in the N-terminal region (right panel). (C) Synonymous codon mutational pattern for mutations with substantially altered ES^CcdB^ as a function of amino acid. Amino acid identity is shown at the circumference of the radar plot. Each circle represents the number of mutants (shown on the vertical axis) obtained for each amino acid, showing a hyperactive (left panel) or inactive (right panel) phenotype.

Rare codons in *E.coli* generally end with A/T, and rare codons ending with A/T are known to correlate with increased expression compared to synonymous mutations ending with G/C (11). This observation is consistent with our results. This association also forms a link to the mRNA transcript secondary structure. If secondary structure is the dominant factor, we would expect a disproportionate enrichment of A over T due to G-U wobble base pairing. GU base pairing is known to be negatively associated with translation efficiency (39). Indeed, nucleotide triplets with A in the wobble positions are enriched in the ‘H+D’ mutant class (Figure 4 B). Disproportionate enrichment of A over T is more prominent at the N-terminus (Figure 4 B). Rare codons in the N-terminal region with increased A/T content likely increase translation initiation, thereby increasing the efficiency with which the gene is translated.

### Arginine displays the maximum phenotypic and codon specific effects

We analysed synonymous mutants giving either a hyperactive phenotype (ES^CcdB^<0.7) or an inactive phenotype (ES^CcdB^>1.8). Synonymous mutations of R, T, S and V are enriched in the former while R alone is enriched in the latter class (Figure 4 C). R and S are encoded by six codons whereas V and T are encoded by four codons.

We also measured the variation of mutational phenotypes amongst the synonymous mutants of the same residue, by analysing the coefficient of variation of ES^CcdB^ scores for each CcdB residue. The coefficient of variation is highest for mutations in arginine, serine, and leucine, suggesting that amino acids with the maximum number of degenerate codons have the largest codon specific effects (Figure S4 A). Of the three, arginine showed the maximum phenotypic effects, depicted by diverse ES^CcdB^ scores (Figure S4 B), perhaps because it encodes part of an anti-SD sequence (AGG) which in turn will modulate the translational rate.

### Single-site synonymous substitutions combined with an N-terminus hyperactive synonymous mutation display enhanced mutational sensitivity

We further incorporated an N-terminal hyperactive synonymous mutation (K4_AAG 285 → K4_AAA) in the background of the existing single-site synonymous mutation library, therefore, generating an exhaustive double-site synonymous mutant library in the presence of a Parent Hyperactive Mutation (PHM). The ES^CcdB^ score of the single-site K4_AAA mutation is 0.42 relative to the K4_AAG WT sequence. The enhanced toxicity implies that a larger amount of CcdB is produced with K4_AAA relative to K4_AAG. This double-site synonymous mutant library was transformed in the three strains as described previously. A good correlation was observed between the two biological replicates in each of the resistant (r= 0.99), sensitive (r= 0.72) and RelE reporter strain (r= 0.99) (Figure 5 A). The number of reads for most mutants in the sensitive strain was very low because of the enhanced toxicity displayed by the K4_AAA mutation due to which correlation between the two biological replicates decreased. Of the ~225 possible mutants, we obtained information for ~160 mutants. Data analysis and assignment of mutant score in the form of E′^CcdB^ and E′^RelE^ for each mutant was done as described previously (32) (Table S1). Here, E′^CcdB^ and E′^RelE^ score for WT K4_AAA is 1. The ‘′’ superscript indicates the scores are associated with the K4_AAA mutant library. There is an issue with the E′^RelE^ scores because ES^RelE^ of the K4_AAA mutation in the WT *ccdB* gene background is 0.35 and is the minimum of the ES^RelE^ values (Table S1). Hence, in the double mutant library it is hard to get ES^RelE^ scores lower than this, a plausible reason why E′^RelE^ scores for most mutants are generally ~1 or >1 (Table S1). Therefore, we do not interpret E′^RelE^ values for the double mutant library. 15 mutants were individually constructed, and E′^CcdB^ scores obtained for these mutants were validated by screening on plates. A good correlation of r = 0.95 was observed between E′^CcdB^ obtained from deep sequencing data and normalised colony count of mutants relative to WT obtained from individually spotting dilutions of mutants transformed in the sensitive strain on plate (Figure 5 B).

**Figure 5:**
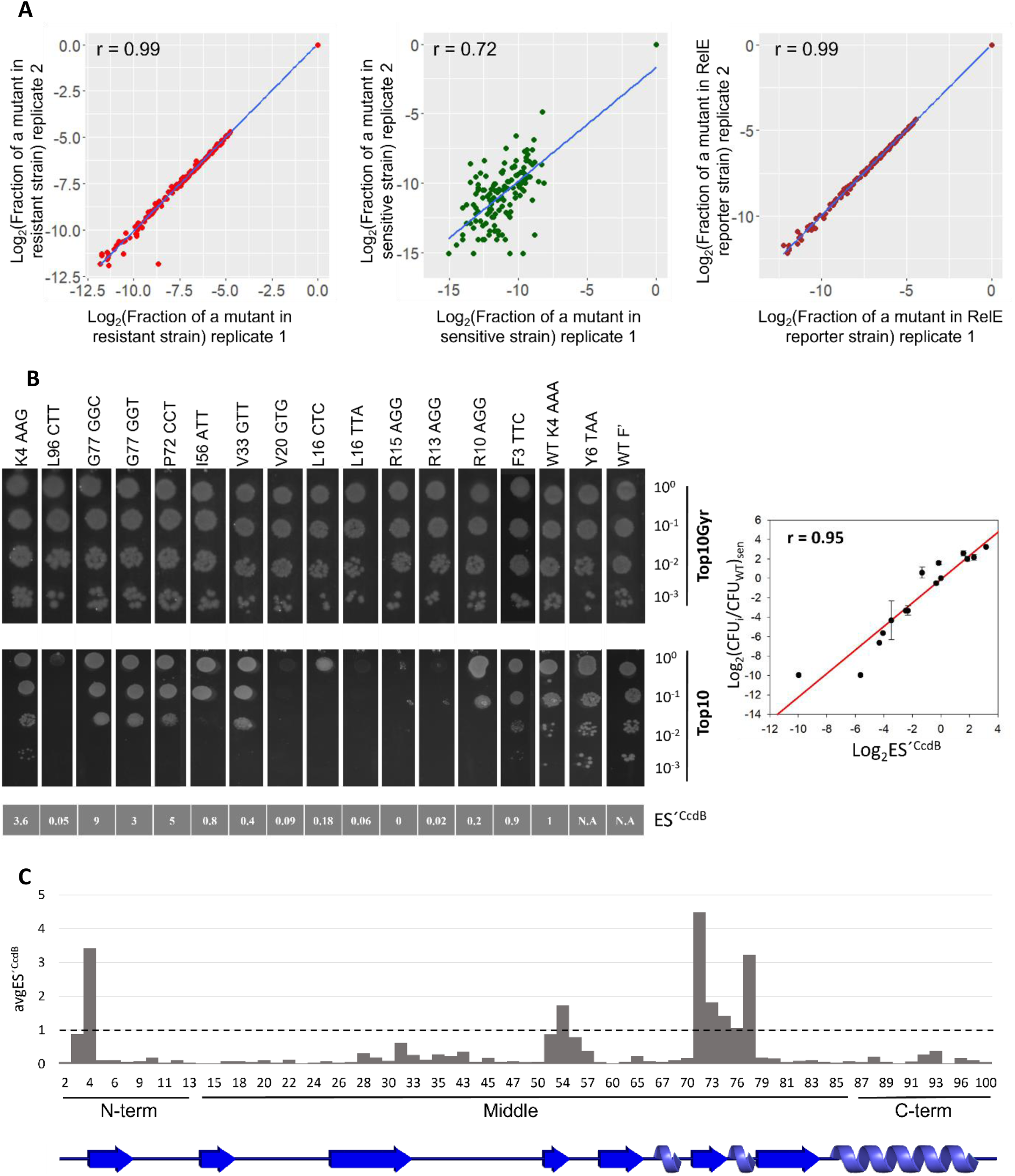
Phenotypic effects of single-site synonymous mutants in the background of an N-terminal hyperactive synonymous mutation, inferred through deep sequencing. (A) Correlations between the biological replicates. The fraction of each mutant normalised to the ‘WT K4_AAA’ value is estimated for all mutants in resistant (extreme left panel), sensitive (middle panel) and RelE reporter strains (extreme right panel). (B) *In vivo* activities of a few individual mutants to validate deep sequencing inferred phenotypes in a low-throughput manner, by transforming individual constructs in resistant (Top10Gyr) and sensitive (Top10) strains (left panel). These mutants are made in the background of WT K4_AAA. E′^CcdB^ scores for individual mutants obtained from deep sequencing are mentioned in the bottom panel. Y6_TAA and WTF’ are used as controls. Y6_TAA is a stop codon mutation at the sixth position of CcdB, thereby making it non-functional. WTF’ is the WT *ccdAB* operon sequence as present in the F plasmid. These two should grow equally well in both resistant and sensitive strains. A correlation of r = 0.95 (*P*<.001) is observed between the E′^CcdB^ scores and normalised CFU count of mutants relative to WT in the sensitive (Top10) strain obtained in the spotting assay. Error bars for the duplicate experiments are also shown (right panel). (C) Effects of double-site synonymous mutations as a function of location in the gene. E′^CcdB^ scores averaged over all mutants for each position. Black dashed line represents the WT K4_AAA score. As done previously, the entire data is divided into N-terminal (1-13), middle (14-86) and C-terminal (87-101) region.

We plotted avgE′^CcdB^ scores for each position, that are obtained after taking an average of all the mutant scores for each position (Figure 5 C). ~82% (127/155) of the mutants displayed E′^CcdB^ scores less than 0.2, i.e., ~8 fold higher toxicity than the WT. We observed that synonymous mutations in specific stretches such as residues 52-57 and 72-77 showed higher values of E′^CcdB^ than synonymous mutations in other locations. Here, in contrast to what was seen for the single-site synonymous mutant library, we observe that mutants lying in the middle region of the gene show the most diverse phenotypes (Figure 5 C).

### Reduced translational rate affects folding kinetics

A possible way by which such synonymous changes can lead either to different final structures or more likely enhanced yield, is by perturbing the protein folding pathway (40, 41). This could occur for example, by a change in translation kinetics which varies as a function of translation pause sites (26). To probe if the synonymous mutations in CcdB could generate such potential ribosomal pause sites, the difference in interaction energies of the ribosome with the anti-SD (aSD) sequence associated with the synonymous mutations relative to WT, were calculated for a window of 10 nucleotides using the RNAsubopt program in the Vienna RNA package (42). A correlation study was conducted separately for the N-terminal (Figure S5 A) and the remaining (middle and C-terminal) region (Figure S5 B). Consistent with the previous observation of the K4_AAG synonymous mutant library, reduced mRNA structure in the N-terminal region increases CcdB translational efficiency, depicted by positive correlation of E′^CcdB^ with ΔGC (r = 0.3, *P* = 0.14) and negative correlation with ΔMFE (r = −0.4, *P* = 0.07). A slower progression of the ribosome at the beginning of the gene is shown by the negative correlation of E′^CcdB^ with ΔaSD (r = −0.4, *P* = 0.11). In the middle and the C-terminal region, in contrast, a weak positive correlation of E′^CcdB^ with ΔaSD (r = 0.1, *P* = 0.35) suggests that translational pausing is accompanied by increased Gyrase binding activity, implying increased yield of properly folded protein likely by a process involving cotranslational folding. Negative correlation of E′^CcdB^ (r = −0.2, *P*<.05) with ΔGC and positive corelation with ΔMFE (r = 0.2, *P* < 0.05) suggest that unstable mRNA results in diminished CcdB activity, possibly because loss of mRNA structure might lead to enhanced *ccdAB* mRNA degradation (Figure 6 A).

**Figure 6:**
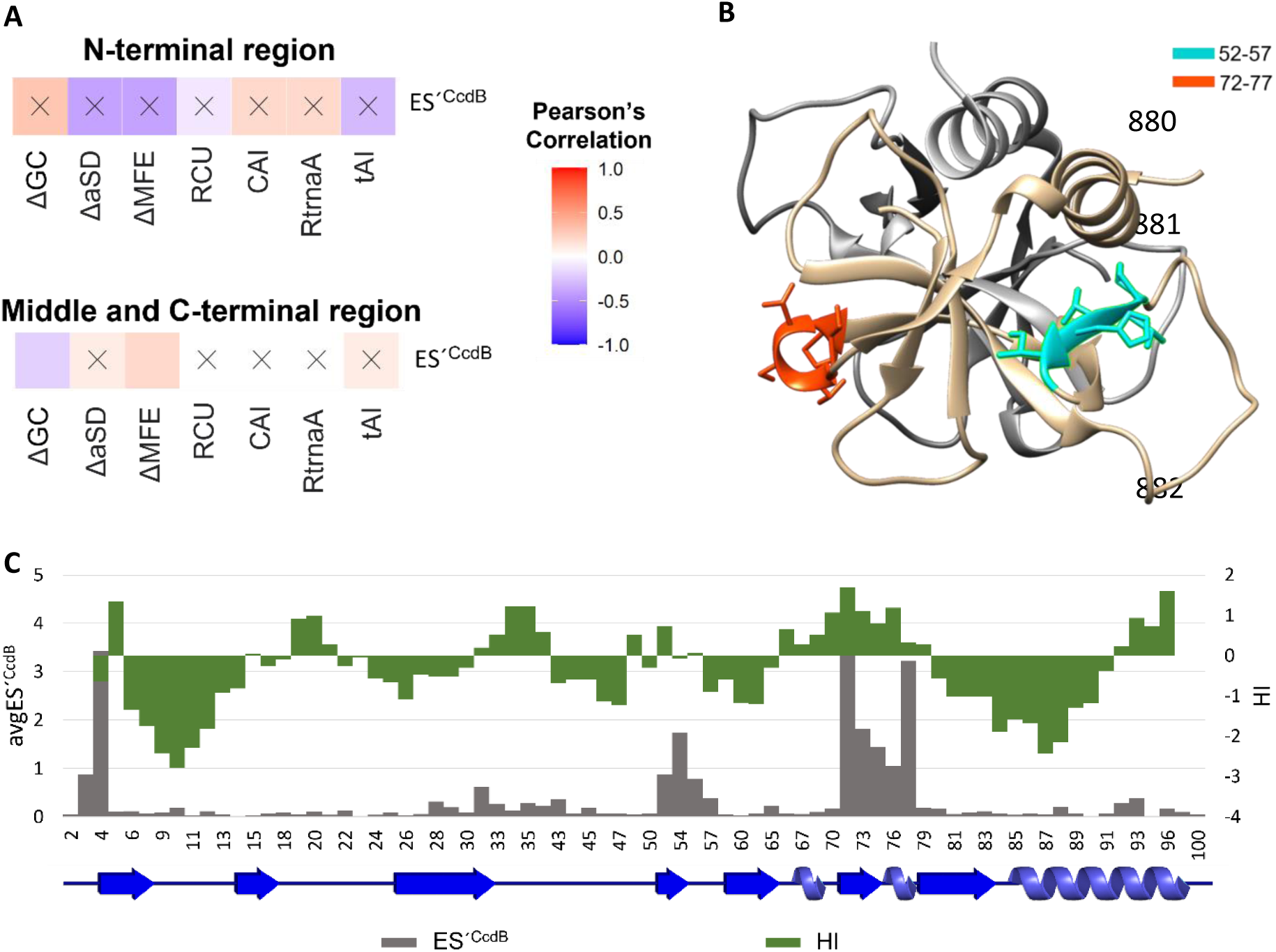
Molecular bases for observed phenotypes. (A) Correlations between experimentally derived mutational scores and sequence-based parameters for the double-site synonymous mutants. Top panel shows these correlations for the N-terminal region. Bottom panel shows these correlations for the middle and C-terminal region. White to red color gradation reflects positive correlation whereas white to blue color gradation reflects negative correlation. Non-significant correlations, i.e., correlations with *P*>0.05 are shown with a cross ‘X’ mark. (B) Elucidating structural properties associated with distinct phenotypic effects. One monomer of CcdB is shown in light grey while residue stretches 52-57 (cyan) and 72-77 (orange) are mapped on the other monomer (tan) of the 3VUB crystal structure. The stretches are part of a β-strand and loop regions. (C) Comparison of the Hydropathy Index (green) and average E′^CcdB^ score (grey) as a function of CcdB position. Hydrophobic residues shown with positive HI values tend to have a higher avgE′^CcdB^ score than the majority of the mutants.

Synonymous mutations in residue stretches 52-57 and 72-77 show a more inactive phenotype compared to the rest of the mutants (Figure 5 C), but do not result in formation of translational pause sites (52-57: avgΔaSD = 0.96, avgRtrnaA = 1.23; 72-77: avgΔaSD = 0.55, avgRtrnaA = 1.4). These residues are part of a β-strand (52–57) and a short helical turn followed by a loop (72-77) respectively (Figure 6 B). The data suggest that the absence of translational pause sites in these regions may cause the protein to misfold inside the cell.

We observed that the generation of aSD sequences and rarer codons due to mutations of residues lying in the α-helical regions (65-67, 73-75 and 84-99) (43) increased CcdB activity (avgE′^CcdB^ = 0.25, avgRCU = 0.86, avgΔaSD = −0.33). It is possible that mutations which reduce translation rate in fast folding structural elements like an α-helix, promote either a more stable protein conformation or a higher yield of properly folded protein. On analysing E′^CcdB^ and Hydropathy Index values for each CcdB residue as a function of position, we observed stretches 52-57 and 72-77 show positive HI values (Figure 6 C), suggesting that hydrophobic regions seem to have special translation kinetic requirements that ensure proper folding of the protein. These analyses suggest lack of translational pausing (high ΔaSD) due to synonymous mutations of hydrophobic residues (HI>0) decreases the amount of properly folded CcdB protein (high E′^CcdB^). Therefore, occurrence of non-optimal codon and internal SD-like sequences in specific regions of the sequence as well as enhanced mRNA stability due to synonymous substitutions reduce the translation rate of the gene, that in turn enhances the yield of folded protein.

## DISCUSSION

In its operonic context, single-site synonymous mutations in *ccdB* toxin display a wide variety of fitness effects. The consistency of our findings with other reports validates our approach to measure the effects of synonymous substitutions. In most prior studies, multiple individual synonymous mutants need to be introduced to see an observable phenotypic effect (11, 20), complicating interpretation of the data. In contrast, in the present *ccd* system, we observe strong phenotypic effects with single synonymous substitutions. Here, we employ two different phenotypic readouts. In the CcdB sensitive Top10 strain, we measure ES^CcdB^ which is a measure of the amount of free CcdB toxin (uncomplexed to CcdA), a higher value of ES^CcdB^ implies a lower amount of free CcdB toxin. In the CcdB resistant strain, Top10GyrA, we measure ES^RelE^, which is a measure of the amount of the CcdA:CcdB complex. A higher value of ES^RelE^ implies a higher amount of complex. Mutational effects are amplified because of the transcriptional autoregulation of the operon and use of a high copy number pUC plasmid.

A thorough understanding of effects of codon bias is central to fields as diverse as biotechnology and molecular evolution. Our results agree with other studies indicating that the initiation phase is the major contributing factor to translational efficiency, followed in order by elongation and the termination phases (44–47). We observe that not just codon rarity but also lack of mRNA structure increases translation initiation (Figure 2 B) (3). Evolutionary conservation also provides insights into factors controlling gene expression and translation (48–50), especially at the N-terminus, likely because translation initiation plays a major role in modulating the rate and efficiency with which the gene will be translated. Although mutations in the N-terminal region show the largest diversity of phenotypes, several synonymous mutations in other regions, result in significant changes in fitness. This suggests that changes in elongation rate can also influence protein yield. This single-site synonymous mutant library when generated in the presence of an N-terminus hyperactive synonymous mutation enhanced mutational sensitivity. It is likely that the N-terminus hyperactive synonymous mutation sufficiently increased translational initiation rate, such that mutational effects in the elongation and termination phase can be more easily observed in this background. For several synonymous mutants, CcdB amounts were increased as seen by decreased E′^CcdB^ scores.

The choice of codons affects translation velocity, which in turn might affect the final conformation or amount of properly folded protein (51, 52). *In silico* analyses revealed that introduction of potential pause sites in the middle of the gene through synonymous mutations, resulted in increased CcdB activity, likely by a process involving co-translational (sequential) folding. Our data suggest that synonymous mutations at different secondary structural elements likely alter translational rate which in turn alters the folding kinetics of the protein. From the results obtained through comparative studies of the mutational scores with Hydropathy Index parameters of the WT gene, we speculate that synonymous mutations at hydrophobic residues disrupt the highly synchronised protein folding process, thereby exposing hydrophobic patches that lead to misfolding, aggregation and lower levels of protein synthesis.

Changes in protein structure and function due to change in translation kinetics and cotranslational folding pathway have been observed for other proteins as well, such as chloramphenicol acetyltransferase (20, 40), suf1 (53), and *Echinococcus granulosus* fatty acid binding protein1 (EgFABP1) (41). Another study reported that a silent mutation of the Ile codon AUC to a rare AUU in the coding sequence of the human MDR1 protein changes translation velocity and affects cotranslational folding. This results in a protein with altered conformation and affinity to its substrates (23). Other studies show slowly translating codon clusters frequently occur at domain boundaries (54, 55), suggesting that translational pausing at rare codons may provide a time delay for optimal sequential folding at defined locations of the nascent polypeptide emerging from the ribosome. A systematic study on protein folding shows that cotranslational folding takes place under quasi-equilibrium conditions, provided translation is slower than folding (56).

We have earlier reported the effects of single-site synonymous mutations on the *ccdA* gene in its native operonic context (35). In that study, synonymous mutations to rarer codons decreased translational efficiency of CcdA eventually leading to more cell death than the WT (35). On the contrary, in the present study we observed synonymous mutations to rarer codons increase CcdB translational efficiency, prominently for the N-terminal region. In both cases variable effects were observed especially at the N-terminus. Due to the smaller length of *ccdA* gene (72 amino acids), it was not divided into three segments to study the effects of synonymous mutations on initiation, elongation and termination phases separately. The study, only examined effects of synonymous mutations in CcdA in an operonic context on cell survival. It was not possible to measure the effects of synonymous mutations on CcdA folding as CcdA is an intrinsically disordered protein. However, in the present study, in addition to CcdB toxicity assay, use of the RelE reporter assay helps clarify molecular mechanisms responsible for phenotypic effects seen for synonymous mutations in CcdB. In addition to highlighting molecular correlates of phenotypic changes associated with synonymous mutations, this study also outlines a novel approach to probe changes in co-translational folding and assembly associated with such single-site synonymous mutations. Such studies help to understand the molecular bases of alteration in gene expression and protein activity arising due to synonymous mutations. It will be interesting to see if such phenotypic effects are observed in other systems. The challenge is to design sensitive readouts wherein small changes in protein activity, result in observable phenotypes. Further focus on development of strategies that can provide direct evidence of transient or permanent perturbations in protein structure arising due to synonymous mutations is also needed.

## MATERIALS AND METHODS

### Plasmids and Host Strains

WT *ccdAB* operon cloned in pUC57 vector (pUCccd) is used as the starting plasmid for making the library of mutants. Two *E. coli* host strains were used, Top10Gyr, containing the GyrA462 mutation resistant to the activity of the CcdB toxin (32), and Top10 which is sensitive to the action of CcdB. These were used for phenotypic screening of CcdB synonymous mutants (32). A third strain is a RelE reporter strain, namely Top10Gyr harbouring a plasmid containing a RelE reporter gene downstream of the *ccd* promoter with a strong Ribosome Binding Site (RBS). The RelE reporter strain is sensitive to the level of the RelE toxin expressed from the *ccd* operon (32).

### Generation of a *ccdB* single-site synonymous mutant library in its native operon

Mutagenic primers for all 100 positions (residues 2 to 101) of *ccdB* were designed such that the degenerate codons were at the 5’ end of each 21 bp forward primer. The entire pUCccd vector backbone was amplified using non-overlapping adjacent 21 bp primers by inverse PCR methodology (34). The primers were obtained in 96-well format from the PAN Oligo facility at Stanford University. A master-mix containing Phusion DNA Polymerase was made for carrying out PCR for all positions in a 96 well format. Following densitometric quantification, an equal amount of PCR product (~200 ng) of each position was pooled. Gel-band purification of the pooled PCR product at the required size (~3.6 Kb) was done using a Fermentas GeneJET™ Gel Extraction Kit according to the manufacturer’s instructions. After purification, pooled PCR product was phosphorylated by T4 PNK, followed by ligation with T4 DNA Ligase. The ligated product was transformed into high efficiency (10^9^ CFU/μg of pUC57 plasmid DNA) electro-competent *E. coli* Top10Gyr cells, and subsequently plated on LB agar plates containing 100 μg/mL ampicillin for selection of transformants. Top10Gyr is referred to as the resistant strain in this study as it is resistant to the toxic activity of the CcdB toxin. Plates were incubated for 12-16 hrs at 37°C. ~100 fold higher number of colonies than the expected library diversity (~200 mutants) were obtained. Pooled plasmid was purified using a Qiagen plasmid maxiprep kit. The entire CcdB synonymous mutant library was then transformed in sensitive (Top10) and RelE reporter strains (Top10Gyr containing the pBT_RelE construct with a consensus SD sequence) in two biological replicates, followed by deep sequencing using the protocol described previously (32). The WT *ccdAB* operon used in this study has a mutation in the putative SD sequence of *ccdA*, which in turn decreases *ccdA* expression, therefore allowing us to screen for both hyperactive and inactive mutants arising from CcdB mutations that confer increased or decreased toxicity relative to WT, respectively. A few mutants were individually made in the same vector, followed by transformation in Top10Gyr and Top10 for low-throughput validation of deep sequencing inferred phenotypes (32).

### Data Normalisation

The raw read numbers for the *ccdB* synonymous mutant library in pUC57 vector were normalised to the total number of reads in each condition. This gave an estimate of the fraction of each mutant represented in that condition. Read numbers for all mutants at all 101 positions (1-101) in CcdB were analysed. Mutants having less than 20 reads in the resistant strain were filtered out prior to subsequent analysis. The rationale for using this read cut-off has been provided previously (35). Two types of variant scores were assigned to each mutant in this study, one is Enrichment Score^CcdB^ (ES^CcdB^) based on the CcdB toxicity readout while the other is Enrichment Score^RelE^ (ES^RelE^) based on the RelE toxicity readout (32).

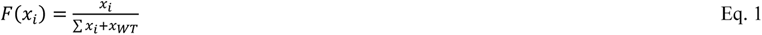

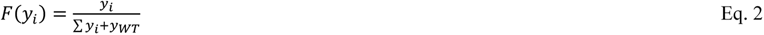

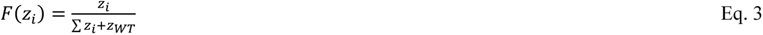

Here, a given mutant is represented by ‘i’ whereas WT is represented by ‘WT’. Number of reads in Top10Gyr resistant strain, Top10 sensitive strain and RelE reporter strain is represented by ‘x’, ‘y’ and ‘z’, respectively. F(x_i_), F(y_i_) and F(z_i_) are the fraction representation of a mutant in resistant, sensitive and RelE reporter strain, respectively.

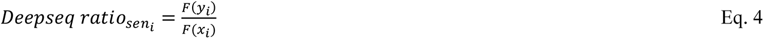

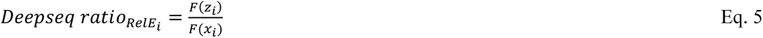

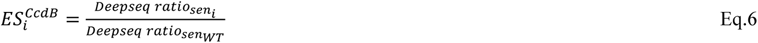

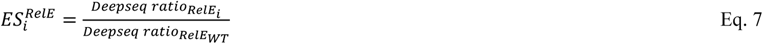

For simplicity, these mutational scores are represented as ES^CcdB^ and ES^RelE^ for each mutant throughout the text. ES^CcdB^ and ES^RelE^ scores for the two biological replicates were estimated. An average of the scores from the two replicates for ES^CcdB^ and ES^RelE^ is taken for downstream analysis. The two variant scores are generally indicated in linear scale throughout the text. WT scores for ES^CcdB^ and ES^RelE^ are 1.

### Defining codon usage parameters

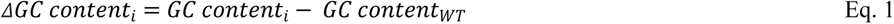

where GC content is the number of Guanine ‘G’ and Cytosine ‘C’ bases present in the mutant codon (GC content_i_) and WT codon (GC content_WT_), respectively.

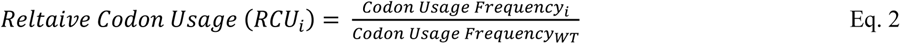

Codon usage frequency values used in this study was taken from Genscript website. These values correlate well (r=0.98) with the codon usage frequency values of *E.coli* K12 strain reported in a study (57). Codon usage frequency is taken from the Bioinformatics tool of Genscript (58). These codon usage values are also mentioned in Table S2.

Codon adaptation index (CAI) is the similarity of codon usage to a reference set of highly expressed genes (59). We calculated CAI for the synonymous mutants of the *ccdB* gene considering *Escherichia coli* (strain K12) as the selected target organism using the Java codon adaptation tool (JCat) (60).

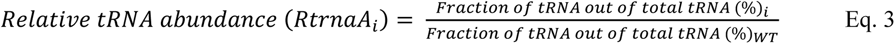

‘Fraction of tRNA out of total tRNA (%)’ for each codon of *E.coli* is the fraction of tRNA out of the total tRNA population in *E. coli* (61). A few degenerate codons are decoded by multiple tRNAs, for simplicity we assume each tRNA binds to its cognate codon with equal probability. Therefore, for such codons we summed the ‘Fraction of tRNA out of total tRNA (%)’ for its corresponding tRNAs and accordingly calculated the RtrnaA_i_.

tRNA Adaptation Index (tAI) is defined as the similarity of codon usage to the relative copy numbers of tRNA genes. tAI computes a weight for each codon, based on the number of tRNAs available in the cell that recognize the codon, and the efficiency of the interaction between the different tRNAs and different codons (62). The score of a coding region is the geometric mean of the weights of all its codons. In order to study the interaction efficiency between a tRNA and a specific codon for the CcdB synonymous mutants, the species-specific tAI (stAI) was calculated using stAI_calc_ (63).

### Defining parameters to estimate the extent to which CcdB levels and codon usage vary relative to WT

Let the CcdB synonymous mutant be denoted by ‘i’ and position ‘j’

If ES^CcdB^_i_<1, then x_i_ = 1/ES^CcdB^

If ES^CcdB^_i_>1, then x_i_ = ES^CcdB^_i_

Let x_minj_ be minimum value of x_i_ for a given position ‘j’

For a given mutant ‘i’ at a particular position ‘j’

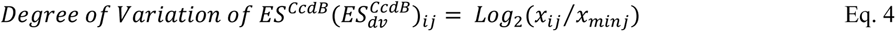

where x_ij_ is the value of x for mutant ‘i’ at position ‘j’.

Similarly, if ES^RelE^_i_<1, then y_i_ = 1/ES^RelE^_i_

If ES^RelE^ >1, then y = ES^RelE^

Let y_minj_ be minimum value of y_i_ for a given position ‘j’

For a given mutant ‘i’ at a particular position ‘j’

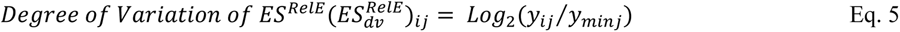

where y_ij_ is the value of y for mutant ‘i’ at position ‘j’.

Similarly, if RCU<1, then z_i_ = 1/RCU_i_

If RCU_i_>1, then z_i_ = RCU_i_

Let z_minj_ be the minimum value of z_i_ for a given position ‘j’

For a given mutant ‘i’ at a particular position ‘j’

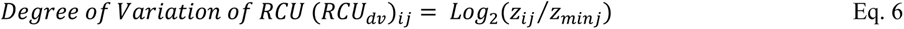

where z_ij_ is the value of z for mutant ‘i’ at position ‘j’.

### Calculation of interaction energies with ribosomal RNA for synonymous mutants in *ccd* mRNA

The interaction energies with the ribosome anti-Shine Dalgarno sequence (5’CACCUCCU 3’) were calculated in the *ccd* mRNA to examine whether the synonymous mutations in the *ccdB* gene region led to generation of SD-like sequences. The difference in the interaction energy with the consensus anti-Shine Dalgarno (aSD) sequence between single synonymous mutants in CcdB and the WT sequence, was calculated for a window of ten nucleotides using the RNAsubopt program from the RNA Vienna package 2.4.18 (42).

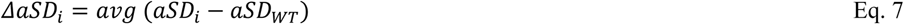

Since aSD values are negative, a positive value of ΔaSD_i_ indicates that the mutant shows lower translational pausing than WT.

### Computational prediction of mRNA secondary structure

A stretch starting 18 bases upstream of the start codon of the transcript, consisting of the putative SD sequence along with the full length 306 bp *ccdB* transcript for WT and all single-site synonymous mutants was used for prediction of secondary structure by the RNAfold program of RNA Vienna package 2.4.18 (42). The entire gene was divided into several segments based on a sliding window of 30 bases. Minimum free energy (MFE) values were computed for each segment, and further averaged over the sliding windows for each mutant. The program calculates the minimum free energy (MFE) structure and outputs the MFE structure and its free energy. We assume the mRNA structure of the toxin is unaffected by the mRNA structure of the preceding antitoxin (28).

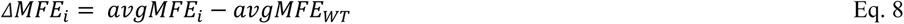

### Nucleotide sequence conservation analysis of *ccdB*

A BlastN search was performed for *WT ccdB* sequence excluding the *E. coli* K12 strain (taxid:83333). The hits were filtered based on ≤ 95% identity. Query coverage was taken between 90 and 100. Sequences were trimmed from the ends such that they were in-frame with the WT sequence. This resulted in 128 sequences which were aligned with CcdB WT sequence using Clustal Omega. This alignment was used as an input for finding the degree of conservation for each position in the CcdB nucleotide sequence using the Consurf web server (http://consurf.tau.ac.il). This is a tool that uses a Bayesian approach to analyse the evolutionary pattern of amino acid and nucleotide sequences (64). The nucleotide preference at each position was further transformed to the preference of codons at the amino acid level.

### Protein sequence conservation analysis of CcdB

Homologous protein sequences of CcdB toxin were identified through the CONSURF server using the HMMER search algorithm for 3 iterations (64). 500 sequences that are part of the list of homologues to query the CcdB sequence, were analysed at a minimum sequence identity of 35%. The maximum sequence identity of non-redundant homologues with each other is less than 95%. Many of these homologues are annotated as CcdB toxins in different organisms. A multiple sequence alignment of the sequences was performed by CONSURF using the MAFFT algorithm, and conserved residues were identified. An evolutionary conservation (Ev cons) score ranging from 1 to 9 (most variable to most conserved) is assigned to each residue.

### Generation of a double site-synonymous mutant library of CcdB in its native operon

Primers were designed to mutate AAG to AAA at the 4^th^ position in the single-site synonymous mutant library via 3-fragment recombination (65) in pUC57 vector. The K4_AAA mutant is considered as the WT gene for this part of the study, which we named as ‘WT K4_AAA’. In this process, other mutations at the first ten positions were lost. Synonymous mutations to all possible degenerate codons were individually generated for these 10 positions via 3-fragment recombination using a Gibson assembly mix. Primers used for amplification of the fragments had 24 bp homology with the gene, and the mutation was present in the middle of the forward primer. These mutants were further pooled with the entire synonymous mutant library in the same ratio as they would be expected to be represented in the library. After synthesising this library and isolating it following transformation in the resistant strain, it was transformed in the sensitive strain and the RelE reporter strain as described previously (32). Following plating, plasmids from each library were isolated, deep sequenced, and assigned variant scores as E′^CcdB^ and E′^RelE^ as described previously (32). After analysis of the data a few individual mutants were selected, synthesised and individually transformed in Top10Gyr and Top10 for low-throughput validation.

## ACKNOWLEDGEMENTS

PB acknowledges University Grants Commission, Government of India, for her fellowship. MB acknowledges Council of Scientific & Industrial Research (CSIR), Government of India, for her fellowship. RV is a J. C. Bose Fellow of DST. This work was funded by grants to RV from the Department of Science and Technology, grant number-EMR/2017/004054, DT.15/12/2018) and Department of Biotechnology, grant no. BT/COE/34/SP15219/2015 DT. 20/11/2015, Government of India. We also acknowledge funding for infrastructural support from the following programs of the Government of India: DST FIST, UGC Centre for Advanced study, Ministry of Human Resource Development (MHRD), and the DBT IISc Partnership Program. The funders had no role in study design, data collection and interpretation, or the decision to submit the work for publication.

## AUTHOR CONTRIBUTION

P.B. and R.V. designed research; P.B. performed research; M.B. assisted with research; P.B., M.B. and R.V. analyzed data; and P.B. and R.V. wrote the paper.

## DATA AVAILABILITY

The raw deep sequencing data used in the present study has been deposited in NCBI’s Sequence Read Archive (accession no. SRR17982061). Rest all study data is included in this article.

