## Supplementary material for "Molecular bases for strong phenotypic effects of single-site synonymous substitutions in the *E.coli ccdB* toxin gene"

### SUPPORTING INFORMATION

#### Supplementary Tables

**Table S1: Mutational scores for synonymous mutants obtained from both K4_AAG and K4_AAA libraries.**

| Mutation | $ES^{CcdB}$ | $ES^{RelE}$ | $ES'^{CcdB}$ | $ES'^{RelE}$ |
|---|---|---|---|---|
| 2Q CAA | 0.06 | 0.59 | 0.05 | 1.40 |
| 3F TTC | 3.29 | 0.69 | 0.88 | 1.45 |
| 4K AAA | 0.42 | 0.35 | - | - |
| 4K AAG | - | - | 3.43 | 1.52 |
| 5V GTC | 0.07 | 0.59 | 0.13 | 1.52 |
| 5V GTA | 0.03 | 0.60 | 0.00 | 1.65 |
| 5V GTG | 0.25 | 0.63 | 0.20 | 1.45 |
| 6Y TAT | 0.07 | 0.77 | 0.12 | 1.47 |
| 7T ACT | 0.04 | 0.67 | 0.03 | 1.38 |
| 7T ACA | 0.04 | 0.89 | 0.09 | 1.31 |
| 7T ACG | 0.02 | 0.66 | 0.08 | 1.45 |
| 8Y TAC | 1.64 | 0.70 | - | - |
| 9K AAG | 1.09 | 0.83 | 0.08 | 1.38 |
| 10R CGT | 3.28 | 0.71 | - | - |
| 10R CGC | 3.83 | 0.64 | - | - |
| 10R CGA | 3.29 | 0.77 | - | - |
| 10R CGG | 2.01 | 0.69 | - | - |
| 10R AGG | 2.79 | 1.04 | 0.19 | 1.42 |
| 11E GAA | 0.21 | 0.99 | 0.04 | 1.41 |
| 12S AGT | 0.21 | 1.00 | 0.05 | 1.31 |
| 12S TCC | - | - | 0.19 | 1.55 |
| 13R CGC | 0.53 | 0.97 | 0.05 | 1.36 |
| 13R CGA | 0.50 | 0.90 | 0.06 | 1.34 |
| 13R CGG | 1.03 | 0.96 | 0.10 | 1.60 |
| 13R AGA | 0.02 | 0.87 | 0.01 | 0.97 |
| 13R AGG | 0.03 | 0.98 | 0.02 | 1.24 |
| 14Y TAC | 0.50 | 1.02 | 0.02 | 1.39 |
| 15R CGC | 0.95 | 1.04 | 0.04 | 1.30 |
| 15R CGA | 0.56 | 1.00 | 0.02 | 1.18 |
| 15R CGG | 0.07 | 0.94 | 0.07 | 1.30 |
| 15R AGA | 1.18 | 0.95 | 0.00 | 1.37 |
| 15R AGG | 0.10 | 0.88 | 0.00 | 1.33 |
| 16L CTT | 0.77 | 1.00 | 0.10 | 1.50 |
| 16L CTC | 0.06 | 0.89 | 0.18 | 1.50 |
| 16L CTA | 1.32 | 1.09 | 0.05 | 1.37 |
| 16L TTA | 1.34 | 1.27 | 0.06 | 1.39 |

| | | | | |
|---|---|---|---|---|
| 16L TTG | 1.04 | 0.37 | 0.02 | 0.86 |
| 17F TTC | 0.74 | 0.77 | - | - |
| 18V GTT | 0.93 | 0.99 | 0.17 | 1.33 |
| 18V GTC | 0.74 | 1.07 | 0.03 | 1.40 |
| 18V GTA | 0.33 | 0.98 | 0.05 | 1.20 |
| 19D GAC | 0.45 | 1.03 | 0.06 | 1.36 |
| 20V GTT | 1.30 | 0.98 | 0.04 | 1.32 |
| 20V GTC | 1.38 | 1.02 | 0.20 | 1.31 |
| 20V GTG | 0.35 | 1.03 | 0.09 | 1.38 |
| 21Q CAA | 1.12 | 0.97 | 0.05 | 1.35 |
| 22S TCT | 0.45 | 0.87 | 0.09 | 1.52 |
| 22S TCC | 1.64 | 1.04 | 0.19 | 1.46 |
| 22S TCA | 0.21 | 1.07 | 0.00 | 1.18 |
| 22S TCG | 0.96 | 0.96 | 0.08 | 1.32 |
| 22S AGC | 1.25 | 1.05 | 0.28 | 1.29 |
| 23D GAC | 1.51 | 1.01 | 0.02 | 1.33 |
| 24I ATC | 0.97 | 1.03 | 0.05 | 1.31 |
| 24I ATA | 0.86 | 0.98 | 0.04 | 1.36 |
| 25I ATC | 0.40 | 1.01 | 0.10 | 1.33 |
| 25I ATA | 0.66 | 1.00 | 0.07 | 1.28 |
| 26D GAT | 1.22 | 0.92 | 0.03 | 1.40 |
| 27T ACT | 0.13 | 0.98 | 0.02 | 1.40 |
| 27T ACC | 0.28 | 1.04 | 0.06 | 1.34 |
| 27T ACA | 0.38 | 1.05 | 0.10 | 1.55 |
| 28P CCT | 0.97 | 1.12 | 0.60 | 1.51 |
| 28P CCA | 0.98 | 1.10 | 0.17 | 0.92 |
| 28P CCG | 0.85 | 1.02 | 0.18 | 1.32 |
| 29G GGT | 0.53 | 0.90 | 0.09 | 1.26 |
| 29G GGC | 1.05 | 1.12 | 0.46 | 1.37 |
| 29G GGA | 1.24 | 1.16 | 0.06 | 1.56 |
| 30R CGT | 0.55 | 1.02 | 0.15 | 1.53 |
| 30R CGG | 2.04 | 1.07 | 0.19 | 1.42 |
| 30R AGA | 0.52 | 0.92 | 0.04 | 1.54 |
| 30R AGG | 0.78 | 0.90 | 0.00 | 1.12 |
| 31R CGT | 0.78 | 1.13 | - | - |
| 31R AGG | 1.05 | 0.80 | 0.62 | 1.83 |
| 33V GTT | 1.08 | 0.97 | 0.41 | 1.40 |
| 33V GTC | 3.03 | 1.04 | 0.24 | 1.31 |
| 33V GTA | 0.90 | 1.05 | 0.17 | 1.14 |
| 34I ATT | 0.91 | 1.00 | 0.13 | 1.37 |
| 34I ATA | 1.94 | 1.02 | 0.13 | 1.29 |
| 35P CCT | 0.80 | 1.23 | - | - |
| 35P CCA | 2.12 | 1.02 | 0.29 | 1.00 |
| 36L CTA | 0.98 | 1.02 | - | - |
| 36L TTG | 1.06 | 0.96 | - | - |
| 37A GCT | 0.83 | 1.13 | - | - |

| | | | | |
|---|---|---|---|---|
| 38S AGC | 0.97 | 1.05 | - | - |
| 39A GCT | 1.67 | 1.02 | 0.32 | 0.89 |
| 39A GCG | 1.12 | 1.17 | - | - |
| 39A GCC | - | - | 0.14 | 1.09 |
| 40R CGA | 0.93 | 0.95 | - | - |
| 41L CTA | 0.58 | 0.73 | - | - |
| 41L TTG | 0.41 | 0.97 | - | - |
| 42L CTA | 0.84 | 0.82 | - | - |
| 42L TTG | 0.93 | 0.72 | - | - |
| 43S TCT | 1.60 | 1.02 | 0.04 | 1.19 |
| 43S TCG | 0.18 | 1.04 | 0.17 | 1.32 |
| 43S TCC | - | - | 0.87 | 1.46 |
| 44D GAC | 0.63 | 0.90 | 0.07 | 1.23 |
| 45K AAG | 0.89 | 0.99 | 0.18 | 1.34 |
| 46V GTT | 0.37 | 1.10 | 0.08 | 1.31 |
| 46V GTA | 1.11 | 1.04 | 0.10 | 1.14 |
| 46V GTG | 2.25 | 1.11 | 0.01 | 1.38 |
| 47S TCT | 0.43 | 1.14 | 0.10 | 1.24 |
| 47S TCA | 1.00 | 1.04 | 0.05 | 1.24 |
| 47S TCG | 1.45 | 1.04 | 0.08 | 1.20 |
| 48R CGC | 1.10 | 1.00 | - | - |
| 48R CGA | 1.09 | 1.11 | - | - |
| 49E GAG | 0.78 | 1.03 | 0.05 | 1.29 |
| 50L CTC | 1.24 | 1.06 | 0.03 | 1.37 |
| 50L CTA | 1.56 | 1.09 | 0.04 | 1.16 |
| 50L CTG | 0.08 | 1.00 | 0.13 | 1.42 |
| 52P CCT | 1.22 | 0.98 | 0.88 | 0.87 |
| 52P CCA | 0.48 | 0.60 | - | - |
| 53V GTT | 0.50 | 1.01 | - | - |
| 54V GTT | 1.21 | 1.05 | - | - |
| 54V GTA | 1.02 | 1.04 | 1.73 | 1.19 |
| 55H CAC | 1.15 | 0.93 | - | - |
| 56I ATT | 0.90 | 1.13 | 0.79 | 0.33 |
| 56I ATA | 0.80 | 0.87 | - | - |
| 57G GGT | 2.03 | 0.99 | 0.39 | 0.75 |
| 57G GGA | 1.02 | 1.20 | - | - |
| 58D GAC | 1.02 | 0.77 | - | - |
| 59E GAG | 0.70 | 0.97 | 0.05 | 1.33 |
| 60S TCT | 0.58 | 1.00 | 0.00 | 0.97 |
| 60S TCC | 0.29 | 0.94 | 0.09 | 1.43 |
| 60S TCA | 1.22 | 0.91 | 0.01 | 1.31 |
| 60S TCG | 1.03 | 1.00 | 0.00 | 1.45 |
| 62R CGT | 1.90 | 0.99 | 0.04 | 1.27 |
| 62R CGA | 1.80 | 1.06 | 0.07 | 1.33 |
| 62R CGG | 1.30 | 0.98 | 0.08 | 1.37 |
| 65T ACT | 0.50 | 0.94 | 0.06 | 1.32 |

| | | | | |
|---|---|---|---|---|
| 65T ACA | 0.46 | 0.90 | 0.17 | 1.19 |
| 65T ACG | 0.34 | 1.03 | 0.46 | 1.35 |
| 66T ACT | 0.45 | 0.92 | 0.12 | 1.27 |
| 66T ACA | 0.36 | 0.94 | 0.08 | 1.31 |
| 66T ACG | 0.53 | 1.06 | 0.03 | 1.36 |
| 67D GAC | 0.78 | 1.09 | 0.06 | 1.35 |
| 69A GCT | 0.93 | 0.99 | 0.13 | 1.29 |
| 69A GCA | 1.08 | 0.95 | 0.09 | 1.30 |
| 69A GCG | 0.98 | 1.02 | 0.08 | 1.27 |
| 70S TCT | 1.22 | 0.91 | 0.00 | 1.20 |
| 70S TCA | 1.58 | 0.94 | 0.12 | 1.24 |
| 70S TCG | 0.44 | 0.87 | 0.02 | 1.02 |
| 70S AGC | 1.15 | 0.95 | - | - |
| 70S TCC | - | - | 0.57 | 1.38 |
| 71V GTA | 1.11 | 0.91 | - | - |
| 72P CCT | 1.96 | 0.88 | 5.31 | 0.33 |
| 72P CCC | 1.67 | 1.21 | - | - |
| 72P CCA | 1.15 | 1.18 | 3.67 | 0.91 |
| 73V GTT | 0.45 | 0.83 | 1.83 | 1.09 |
| 74S TCT | 1.09 | 1.60 | - | - |
| 74S TCA | 0.75 | 0.99 | 1.44 | 1.89 |
| 75V GTC | 0.87 | 0.83 | - | - |
| 76I ATT | 0.99 | 1.14 | 1.27 | 0.26 |
| 76I ATA | 1.18 | 0.88 | 0.85 | 0.19 |
| 77G GGT | 1.17 | 0.97 | 3.23 | 0.72 |
| 77G GGC | 0.61 | 1.01 | - | - |
| 77G GGA | 1.10 | 1.09 | - | - |
| 78E GAG | 1.03 | 0.71 | - | - |
| 79E GAG | 0.79 | 0.80 | 0.19 | 1.04 |
| 80V GTT | 0.95 | 1.01 | 0.19 | 1.25 |
| 80V GTC | 1.72 | 0.97 | 0.13 | 1.30 |
| 80V GTA | 0.50 | 1.01 | 0.17 | 1.30 |
| 81A GCC | 0.89 | 0.89 | 0.04 | 1.27 |
| 81A GCA | 0.69 | 0.89 | 0.06 | 1.24 |
| 81A GCG | 0.72 | 0.90 | 0.07 | 1.15 |
| 82D GAC | 0.90 | 1.02 | 0.10 | 1.26 |
| 83L CTT | 0.72 | 0.95 | 0.15 | 1.31 |
| 83L CTA | 0.75 | 0.96 | 0.07 | 1.33 |
| 83L CTG | 0.84 | 0.96 | 0.13 | 1.27 |
| 84S TCT | 0.42 | 0.98 | 0.06 | 1.26 |
| 84S TCC | 0.44 | 1.03 | 0.05 | 1.27 |
| 84S TCA | 0.51 | 1.08 | 0.09 | 1.23 |
| 84S TCG | 0.51 | 0.95 | 0.07 | 1.31 |
| 85H CAT | 0.91 | 1.01 | 0.04 | 1.19 |
| 86R CGT | 1.05 | 1.00 | 0.14 | 1.10 |
| 86R CGA | 0.89 | 1.00 | 0.06 | 1.30 |

| | | | | |
|---|---|---|---|---|
| 86R CGG | 0.95 | 0.89 | 0.04 | 1.40 |
| 87E GAG | 0.83 | 1.06 | 0.05 | 1.18 |
| 88N AAC | 0.95 | 0.96 | 0.21 | 1.28 |
| 89D GAT | 0.97 | 0.96 | 0.06 | 1.20 |
| 90I ATT | 0.97 | 1.04 | 0.02 | 1.16 |
| 90I ATA | 1.05 | 0.96 | 0.03 | 1.33 |
| 91K AAG | 0.71 | 0.55 | 0.07 | 1.13 |
| 92N AAT | 1.40 | 0.95 | 0.28 | 1.25 |
| 93A GCT | 0.98 | 1.02 | - | - |
| 93A GCA | 0.79 | 1.20 | 0.38 | 1.14 |
| 94I ATC | 1.19 | 1.01 | - | - |
| 94I ATA | 0.84 | 0.76 | - | - |
| 95N AAT | 1.00 | 1.02 | 0.02 | 1.24 |
| 96L CTT | 1.17 | 1.05 | 0.05 | 1.32 |
| 96L CTC | 1.13 | 1.03 | 0.06 | 1.26 |
| 96L CTA | 1.32 | 0.96 | 0.38 | 1.35 |
| 98F TTT | 0.94 | 0.86 | 0.10 | 1.28 |
| 100G GGT | 0.34 | 1.00 | 0.05 | 1.21 |
| 100G GGC | 1.14 | 1.02 | 0.04 | 1.30 |
| 100G GGG | 0.93 | 0.97 | 0.07 | 1.34 |

**Table S2: Normalized codon usage for different degenerate codons encoding the same amino acid.**

| Codon | Amino acid | Fraction[a] | Normalised codon usage[b] |
|---|---|---|---|
| GCT | A | 0.18 | 0.00 |
| GCC | A | 0.26 | 0.53 |
| GCA | A | 0.23 | 0.33 |
| GCG | A | 0.33 | 1.00 |
| TGT | C | 0.46 | 0.00 |
| TGC | C | 0.54 | 1.00 |
| GAT | D | 0.63 | 1.00 |
| GAC | D | 0.37 | 0.00 |
| GAA | E | 0.68 | 1.00 |
| GAG | E | 0.32 | 0.00 |
| TTT | F | 0.58 | 1.00 |
| TTC | F | 0.42 | 0.00 |
| GGT | G | 0.35 | 0.92 |
| GGC | G | 0.37 | 1.00 |
| GGA | G | 0.13 | 0.00 |
| GGG | G | 0.15 | 0.08 |
| CAT | H | 0.57 | 1.00 |
| CAC | H | 0.43 | 0.00 |
| ATT | I | 0.49 | 1.00 |
| ATC | I | 0.39 | 0.74 |
| ATA | I | 0.11 | 0.00 |
| AAA | K | 0.74 | 1.00 |
| AAG | K | 0.26 | 0.00 |
| TTA | L | 0.14 | 0.23 |
| TTG | L | 0.13 | 0.21 |
| CTC | L | 0.1 | 0.14 |
| CTA | L | 0.04 | 0.00 |
| CTG | L | 0.47 | 1.00 |
| CTT | L | 0.12 | 0.19 |
| ATG | M | 1 | 1.00 |
| AAT | N | 0.49 | 0.00 |
| AAC | N | 0.51 | 1.00 |
| CCT | P | 0.18 | 0.14 |
| CCC | P | 0.13 | 0.00 |
| CCA | P | 0.2 | 0.19 |
| CCG | P | 0.49 | 1.00 |

| | | | |
|---|---|---|---|
| CAA | Q | 0.34 | 0.00 |
| CAG | Q | 0.66 | 1.00 |
| CGT | R | 0.36 | 1.00 |
| CGC | R | 0.36 | 1.00 |
| CGA | R | 0.07 | 0.09 |
| CGG | R | 0.11 | 0.22 |
| AGA | R | 0.07 | 0.09 |
| AGG | R | 0.04 | 0.00 |
| TCT | S | 0.17 | 0.27 |
| TCC | S | 0.15 | 0.09 |
| TCA | S | 0.14 | 0.00 |
| TCG | S | 0.14 | 0.00 |
| AGT | S | 0.16 | 0.18 |
| AGC | S | 0.25 | 1.00 |
| ACT | T | 0.19 | 0.09 |
| ACC | T | 0.4 | 1.00 |
| ACA | T | 0.17 | 0.00 |
| ACG | T | 0.25 | 0.35 |
| GTT | V | 0.28 | 0.61 |
| GTC | V | 0.2 | 0.17 |
| GTA | V | 0.17 | 0.00 |
| GTG | V | 0.35 | 1.00 |
| TGG | W | 1 | 1.00 |
| TAT | Y | 0.59 | 1.00 |
| TAC | Y | 0.41 | 0.00 |

NOTE: [a]Codon usage frequency for degenerate codons for the same amino acid for *E.coli* is taken from the Bioinformatics tool of Genscript (58). [b]Normalised Codon Usage signifies codon usage frequency value normalised between 0 and 1 where 0 signifies the rarest codon and 1 signifies the most preferred codon for a particular amino acid.

#### Supplementary Figures

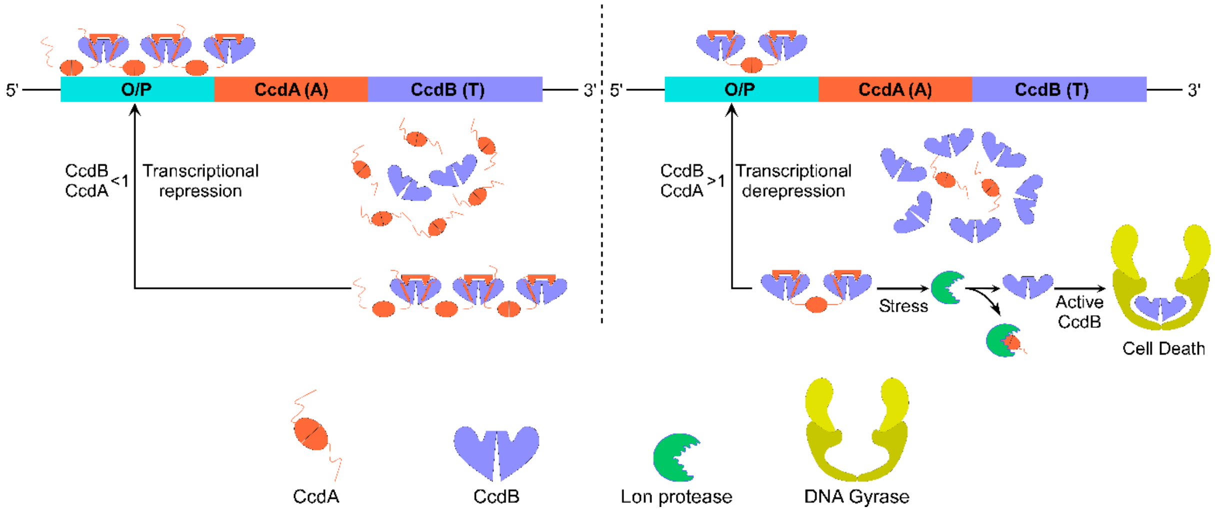


**Figure S1: Schematic of the autoregulation of the CcdAB complex.** Autoregulation is a key feature of Toxin-Antitoxin systems that determines the toxin:antitoxin ratio within cells. The left panel shows that when $[CcdA]_{TOT} > [CcdB]_{TOT}$, the CcdA-CcdB proteins form an extended multimeric complex that represses transcription of the operon. CcdA alone binds weakly to the O/P region. CcdB binding to CcdA allows CcdA to bind tightly to the O/P region. The right panel shows that when $[CcdA]_{TOT} < [CcdB]_{TOT}$, the CcdA-CcdB proteins form a CcdB:$(CcdA)_2$:CcdB heterotetramer which binds weakly to the O/P region and derepresses the transcription of the operon.

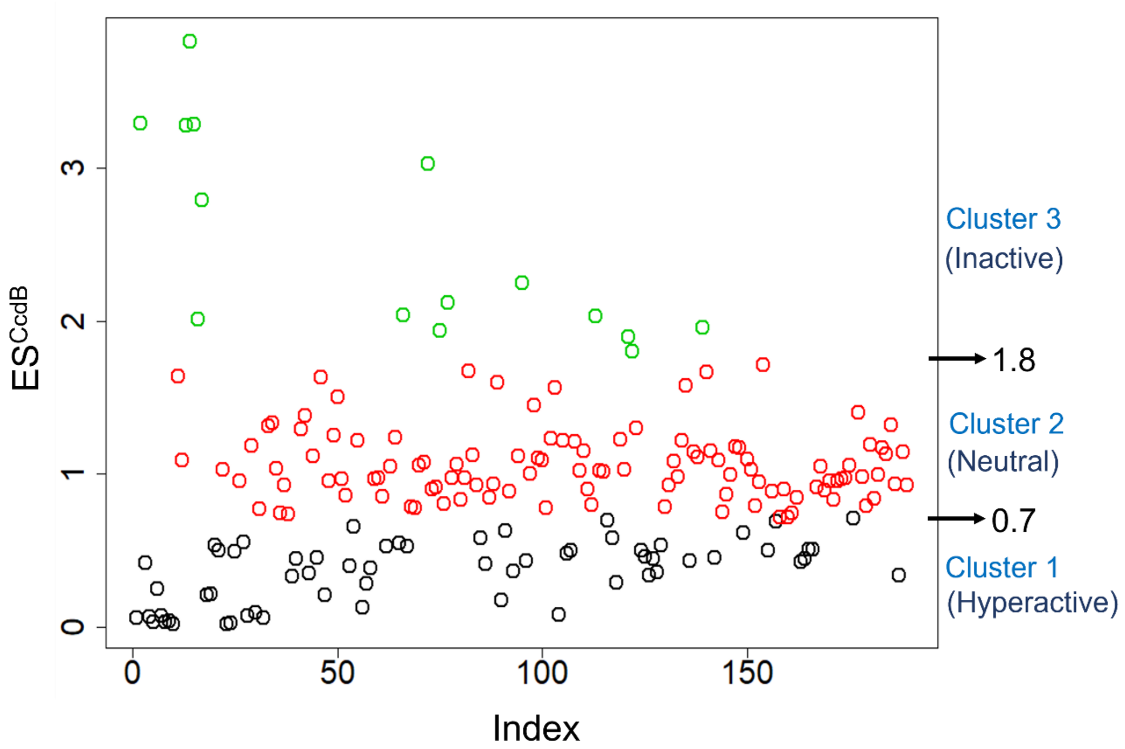


**Figure S2: Division of CcdB synonymous mutants into three classes based on ES$^{CcdB}$ values.** K-means clustering divides ES$^{CcdB}$ values for synonymous mutants into three clusters. Green, red, and black dots denote inactive, neutral and hyperactive phenotypes, respectively. Cluster boundaries are denoted by horizontal arrows. These values are used to classify mutants as hyperactive (ES$^{CcdB}$<0.7), neutral (0.7<ES$^{CcdB}$<1.8) and inactive (ES$^{CcdB}$>1.8) respectively. Cluster 1, 2 and 3 contain 62, 112 and 15 synonymous mutants, respectively. Mean values of cluster 1, 2 and 3 are 0.363, 1.06 and 2.5, respectively.

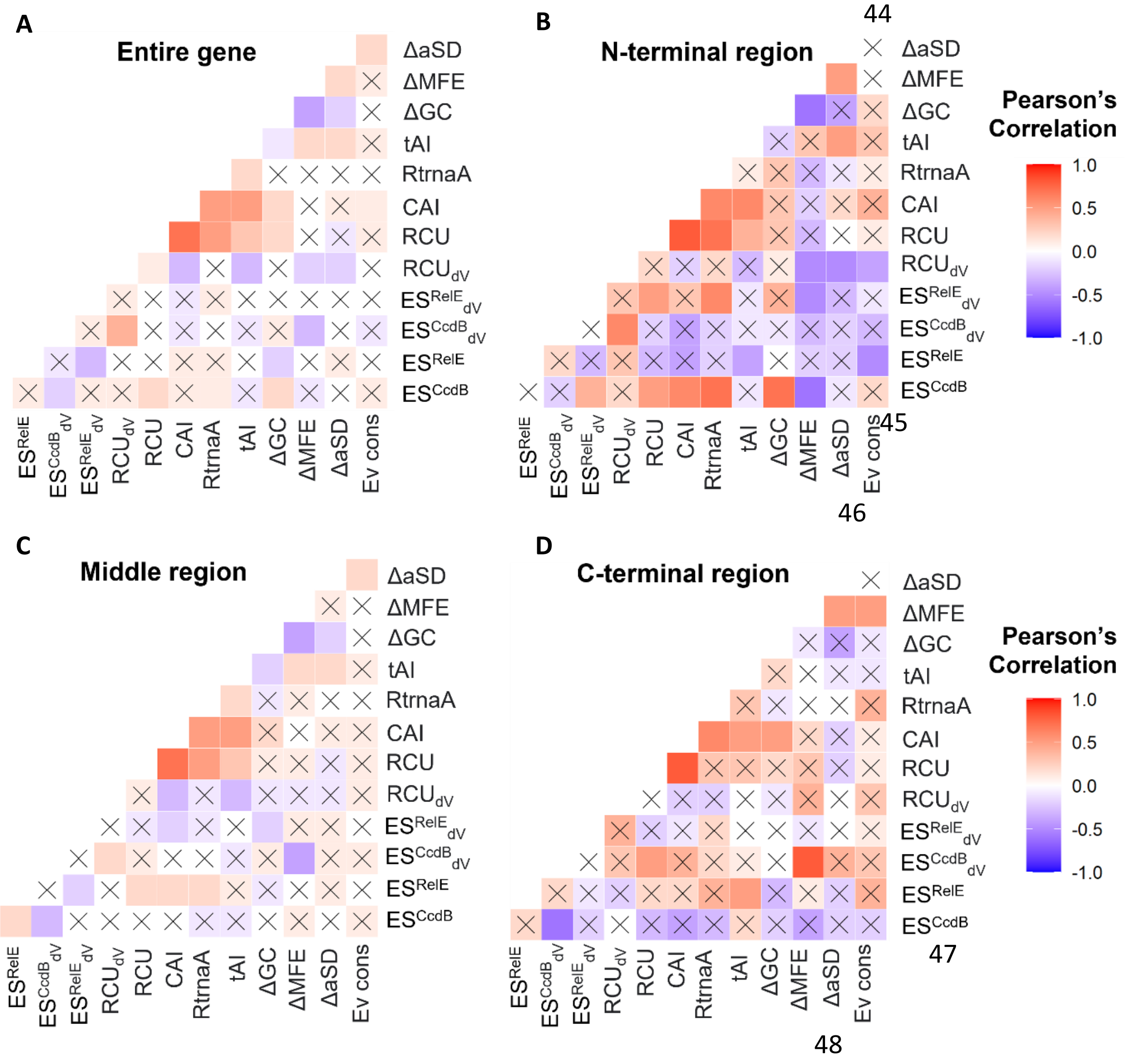


**Figure S3: Insights into the molecular bases for the observed phenotypic effects for the single-site synonymous mutant library.** (A-D) Correlation between experimentally determined relative fitness values and various sequence-based parameters for (A) Entire gene, (B) N-terminal, (C) Middle and (D) C-terminal regions. Colour bar for the Pearson's correlation coefficient is shown on extreme right. Correlations with $P>0.05$ are marked with a cross 'X'. Parameter definitions are in the Materials and Methods section.

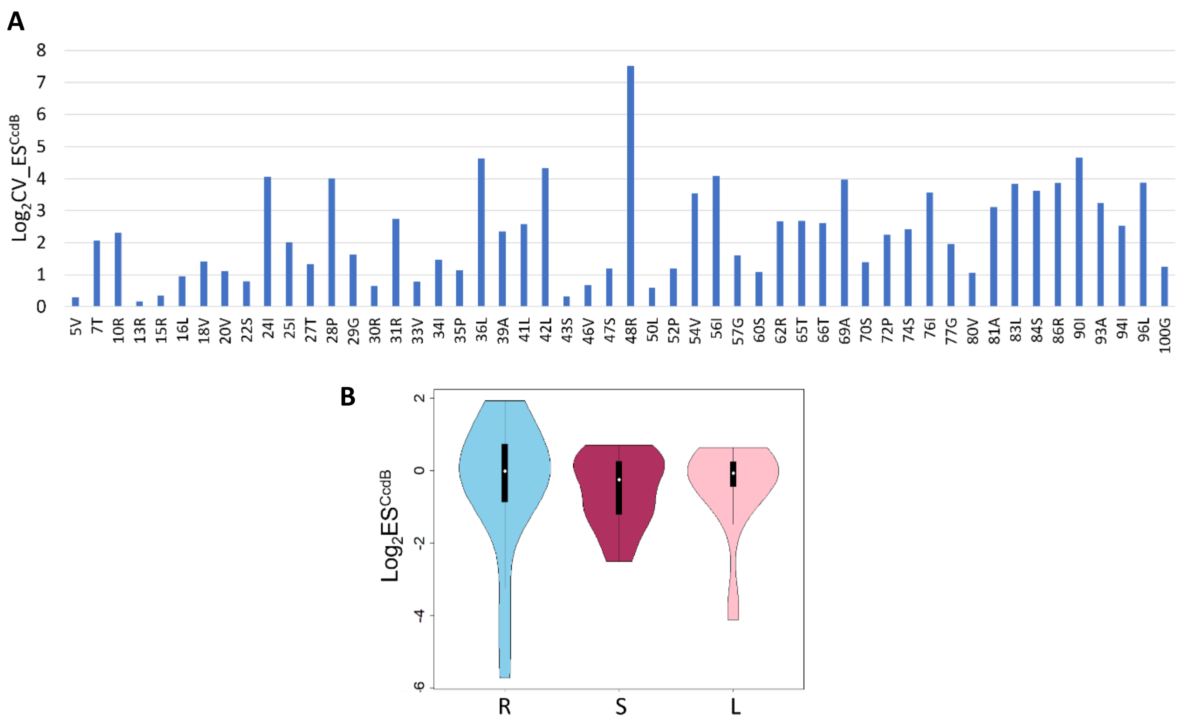


**Figure S4: Distributions of codon-specific effects for the synonymous mutant library.** (A) Coefficient of variation (CV) of $ES^{CcdB}$ score for each residue with two or more codons is plotted as a function of residue number with its corresponding WT amino acid indicated. Codon-specific effects are measured by calculating the coefficient of variation (mean/s.d.) of $ES^{CcdB}$ for each mutant residue with two or more mutants in the synonymous mutant library. (B) Distribution of $ES^{CcdB}$ for amino acids with six degenerate codons. Arginine displays the largest diversity in phenotypic effects.

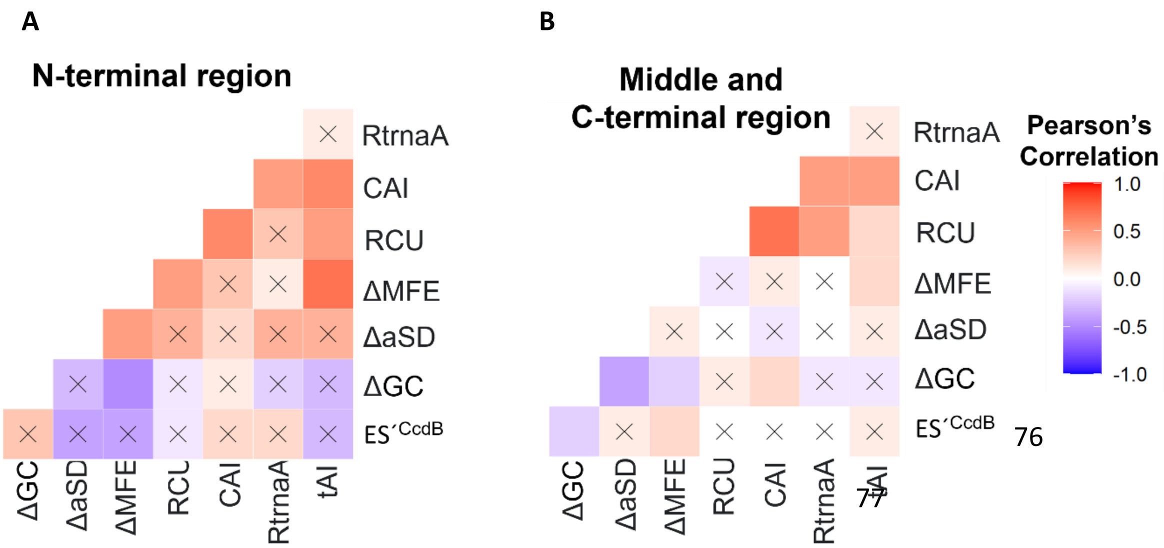


**Figure S5: Insights into the molecular bases for the observed phenotypic effects of the double-site synonymous mutant library, i.e., in background of K4_AAA mutation.** Correlation between experimentally determined relative fitness values and various sequence-based parameters for (A) N-terminal (Residue no. 1-13) and (B) Middle and C-terminal (Residue no. 14-101) regions. Colour bar for the Pearson's correlation coefficient is shown on extreme right. Correlations with *P*>0.05 are marked with a cross 'X'. Parameter definitions are in the Materials and Methods section.
